# Neuroimaging model of visceral manipulation in awake rat

**DOI:** 10.1101/2024.09.17.613477

**Authors:** Samuel R. Cramer, Xu Han, Dennis C.Y. Chan, Thomas Neuberger, Nanyin Zhang

## Abstract

Reciprocal neuronal connections exist between the internal organs of the body and the nervous system. These projections to and from the viscera play an essential role in maintaining and finetuning organ responses in order to sustain homeostasis and allostasis. Functional maps of brain regions participating in this bidirectional communication have been previously studied in awake humans and anesthetized rodents. To further refine the mechanistic understanding of visceral influence on brain states, however, new paradigms that allow for more invasive, and ultimately more informative, measurements and perturbations must be explored. Further, such paradigms should prioritize human translatability. In the current paper, we address these issues by demonstrating the feasibility of non-anesthetized animal imaging during visceral manipulation. More specifically, we used a barostat interfaced with an implanted gastric balloon to cyclically induce distension of a non-anesthetized rat’s stomach during simultaneous BOLD fMRI. General linear modeling and spatial independent component analysis revealed several regions with BOLD activation temporally coincident with the gastric distension stimulus. The ON-OFF (20 mmHg - 0 mmHg) barostat-balloon pressure cycle resulted in widespread BOLD activation of the inferior colliculus, cerebellum, ventral midbrain, and a variety of hippocampal structures. These results suggest that neuroimaging models of gastric manipulation in the non-anesthetized rat are achievable and provide an avenue for more comprehensive studies involving the integration of other neuroscience techniques like electrophysiology.

**Significance Statement:** It is unclear to what extent measurements of brain activity are affected by background, and experimentally unrelated, interoceptive processes. To advance our understanding of ongoing visceral activity’s influence on brain states, here we provide a proof of concept, anesthesia-free animal model of visceral manipulation during simultaneous BOLD fMRI. We successfully demonstrated BOLD activation during gastric distension of the unanesthetized rat in both classically reported (cerebellum, hippocampus) and novel (inferior colliculus) regions. This paradigm establishes an important foundation for further interrogation of viscera-brain interactions.

## 1. INTRODUCTION

Neuronal tracing studies have firmly established the existence of bidirectional (afferent and efferent) communications between the brain and viscera (internal organs) ^1–4^. These afferent and efferent pathways provide precise moment-to-moment monitoring and regulation, respectively, of the viscera. This fast-paced, life-essential dynamic suggests a fundamental relationship between visceral states and brain activity. However, the prevailing models of brain structures influenced by or participating in these visceral states remains incomplete.

To better understand the role of visceral states in brain activation patterns, it is important to first establish a model that 1) manipulates viscera to reliably induce altered brain activity 2) maximizes the spatial coverage of measured brain responses. Regarding the first criterion, the stomach is an excellent candidate organ to manipulate due to its comparatively ease of access, structural resilience, and large number of neurons involved in communication with the brain. Regarding the second criterion, both positron emission tomography (PET) and magnetic resonance imaging (MRI) techniques enable measurements with considerable spatial coverage. Unsurprisingly, at least two decades of prior research has provided an initial framework for such a model. For example, H_2_O^15^- PET research from early 2000’s demonstrated in humans that block design intragastric balloon inflation paradigms could be used to induce changes in the brain. These experimenter-controlled distensions of the stomach were temporally coincident with cerebral blood flow (CBF) increases in a variety of locations: cerebellum, insula, anterior cingulate gyrus, thalamus, primary and secondary sensory cortices, and various segments of the brain stem ^5–9^. Additionally, similarly designed sensory-evoked, human fMRI intragastric balloon studies showed blood-oxygen-level-dependent (BOLD) activations in a largely overlapping set of regions ^10–13^. Of course, all such human paradigms are limited due to ethical concerns: uncomfortable oral insertions of balloons through the esophagus prevent large numbers of per-subject trials and invasiveness of direct electrophysiology of the gastric wall prevents relevant measurements of gastric state, to name a few. These limitations have motivated the pursuit of similarly themed experiments in animals, where such considerations are relaxed. Several gastric distension fMRI studies in the rat have been published in the last decade, reporting increased BOLD responses in the hypothalamus, thalamus, nucleus tractus solitarius (NST), hippocampus, cingulate cortex, cerebellum, and the insular and sensory cortices ^14,15^. Unfortunately, *all* rodent experiments to date have been conducted under the influence of anesthesia. Given anesthesia’s disturbance of *normal* sensory information processing ^16–20^, as well as the large list of known anesthesia-fMRI confounds ^21^, non-anesthetized animal imaging serves as an ideal alternative with the promise of greater translatability. In the current paper, unanesthetized animal fMRI was used to investigate the brain effects of gastric distension in the rat. Using a block-design paradigm, we demonstrate that gastric distension of the proximal stomach of an unanesthetized rat is temporally correlated with the BOLD activation of the inferior colliculus, cerebellum, ventral midbrain, and a variety of hippocampal structures.

## 2. METHODS

### 2.1. Animals

All procedures were carried out under a protocol approved by the Pennsylvania State University Institutional Animal Care and Use Committee (IACUC). Male Long Evans rats (Charles River, Wilmington, MA) were used throughout this project (n=19). Upon arriving at the animal facility, all rodents were given at least 5 days of habituation to their new environment (single-housed caging) prior to the start of the overall experimental paradigm. At the time of the first procedure, rats’ ages and weights ranged from 3-4 months and 400g-500g, respectively. Standard rat chow and water were provided ad libitum unless explicitly stated otherwise (refer to Methods 2.3 and 2.4). Room temperature was kept at 22–24 °C with a 12 h light, 12 h dark cycle.

### 2.2. Non-anesthetized animal imaging acclimation

This project involved BOLD fMRI data acquisition of animals in the non-anesthetized condition. Such data collection requires that animals tolerate a loud environment (110 dB+) while undergoing body restraint. Further, animals must demonstrate minimal head motion throughout the imaging procedure. To ensure low-motion profiles across subjects, animals in this experiment were first subjected to non-anesthetized imaging acclimation across a 7-day period. As outlined in Dopfel et al. (2018), such procedures can be used to familiarize non-anesthetized animals to the MRI environment in preparation for eventual data collection ^22^. Here a modified approach was taken to better identify subjects that exhibit desirable motion criteria. Across all seven days, during a 20-to-30-minute exposure to low-dose isoflurane (∼1% conc.), animals were first situated in a custom, in-house-designed, 3D-printed restrainer (Figure 1A). After completing restrainer setup, isoflurane administration was discontinued, allowing animals to regain consciousness. On days 1-4, after waking up from anesthesia, animals were placed inside a mock-MRI environment (non-illuminated, acoustically-insulated box) where prerecorded gradient-echo echo planar imaging (GE-EPI) pulse sequences played at increasing sound intensities across days: Day 1 = 100 dB, Day 2 = 103 dB, Day 3 = 106 dB, and Day 4 = 108 dB. Across these four days, the duration of sound exposure also progressively increased: Day 1 = 15 minutes, Day 2 = 30 minutes, Day 3 = 45 minutes, and Day 4 = 60 minutes. On each of the remaining three days (Day 5, Day 6, and Day 7), after waking from anesthesia, animals were, instead, placed inside a 7T MRI for an imaging session comprised of four 10-minute single-shot GE-EPI scans (more detailed MRI descriptions are offered in Methods 2.4). Prior to each imaging session, an animal’s non-anesthetized status was confirmed via real-time respiration monitoring (Small Animal Instruments Inc.). On each of the seven days, at the conclusion of each faux-imaging / real-imaging session, animals were removed from the mock-MRI environment / 7T MRI, briefly anesthetized for restrainer removal, and returned to their home-cages. Because the final three days of acclimation were performed in a real scanner, head motion data could be quantitatively assessed. Animals with acceptable levels of motion were moved on to the next phase of the project (more details offered in Methods 2.5); animals that did not meet the motion criteria were removed from the study. All remaining animals were given at least seven days of rest between the conclusion of acclimation and the start of surgery.

**Figure 1:**
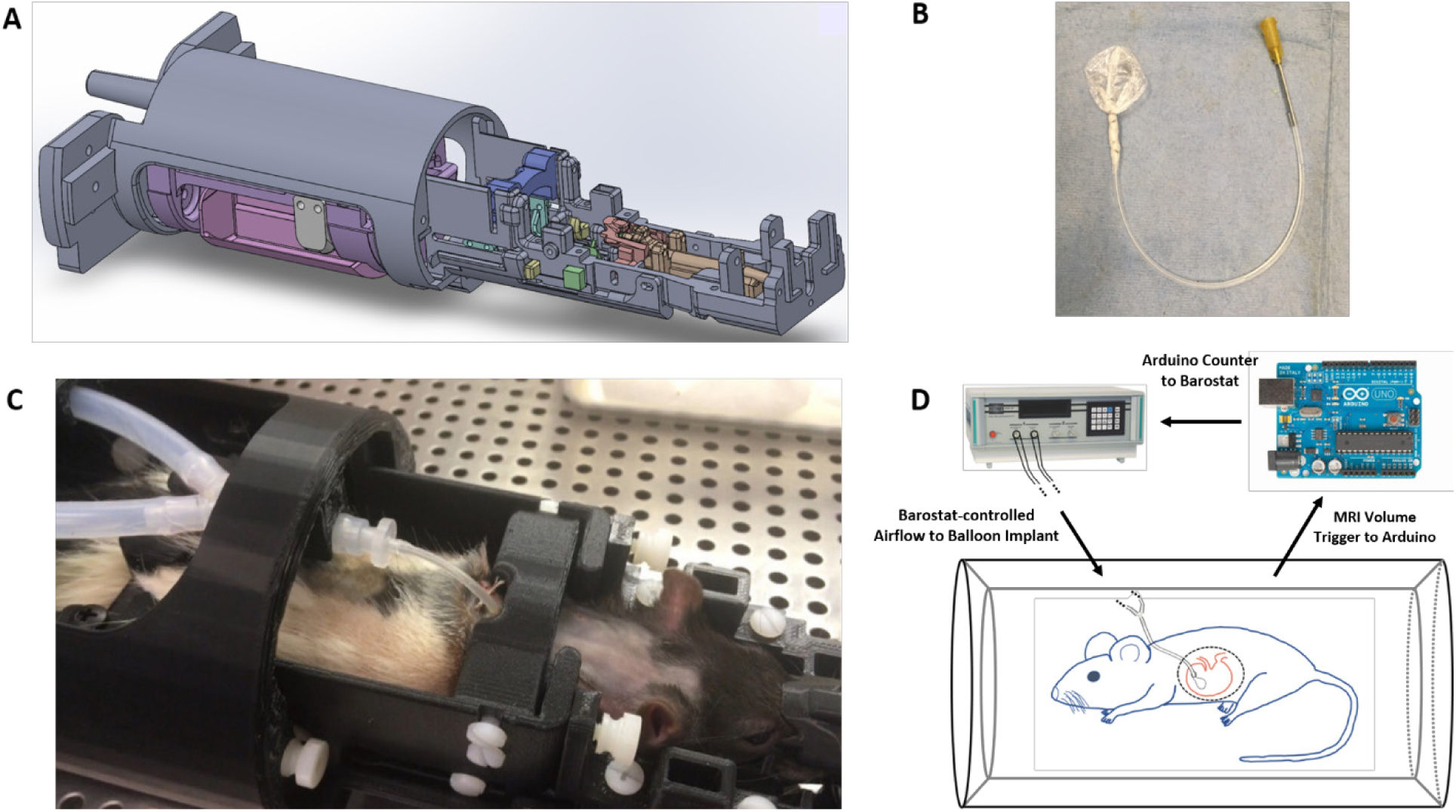
Equipment used in simultaneous fMRI – gastric distension experiment. **(A)** CAD software (SolidWorks) rendering of the complete restraining device used in non-anesthetized rat fMRI. **(B)** Photo depicting the gastric implant. The balloon portion of the implant is comprised of polyethylene and measures approximately 25 mm in diameter; the base of the balloon is attached to a silicon catheter via multiple knots (nylon suture) and parafilm. **(C)** Photo capturing the imaging-specific restrainer, which uses a Y-connecter to interface the scapula-exiting catheter with the barostat tubing. **(D)** Illustration of the tripartite system that allows for proper synchronization between the barostat pressure sequence and the MRI data acquisition sequence.

### 2.3. Gastric balloon implantation surgery

Animals were fasted for 16-18 hours prior to aseptic gastric balloon implantation surgery to minimize food content in the stomach. The gastric balloon implant (Figure 1B) was made in-house using a patch of low-density polyethylene (Gorilla Supply Gloves), a silicone rubber catheter (Liveo^TM^ Pharma-50 Tubing) with a 1.02 mm ID and 2.16 mm OD, nylon filament, and parafilm. When flattened out, the approximately circular object measured ∼25 mm in a diameter. The polyethylene provided a sufficiently non-compliant (for the experiment-related pressures used) material with a maximal liquid volume of ∼5 mL of incompressible water before irreversible deformation of the balloon wall took place. Visual assessment during surgery confirmed that, upon inflation, a balloon of this size was consistently in contact with some portion of the stomach wall. Additionally, the minimized material compliance ensured that the balloon’s volume was predominantly restricted to the fundus and in minimal contact with the proximal portion of the stomach’s corpus/body domain. Prior to surgery, the implant was sterilized with ethylene-oxide. At the time of surgery, animals were anesthetized with low-dose isoflurane, and subcutaneous injections of warm saline (10 mL/kg), Baytril^®^ (2.5 mg active compound/kg), and long acting buprenorphine (1 mg/kg) were administered. Throughout the duration of the surgery, various physiological parameters (heart rate, SpO_2_, and body temperature) were measured and maintained (Kent Scientific Corporation). The stomach was accessed through the abdominal cavity, and a small incision was placed at the pole of the gastric fundus. The implant was inserted into the incision such that the entire balloon was within the stomach; only a small amount (< 2 mm) of a parafilm-wrapped section of the catheter entered the incision. After insertion, the fundus’ incision was sealed, and the catheter was anchored to the fundus wall at the incision site, which ensured that the balloon remained a fixed distance away from the incision. The catheter was passed through, and anchored against, the left lateral abdomen. The catheter was then tunneled subcutaneously around the animal’s side and exited through the skin overlaying the scapula, with a third anchoring suture being placed at the scapular exit site. At the conclusion of the surgery, an empty syringe was used to confirm that the catheter remained patent and allowed for air-driven balloon inflation. Animals were given at least seven days of recovery before moving on to the next phase of the project (Figure 2A). Post-operation treatment included two days of exclusively wet food (DietGel^®^) and seven days of orally-delivered antibiotic treatment through Baytril^®^-treated water (0.7% solution).

**Figure 2:**
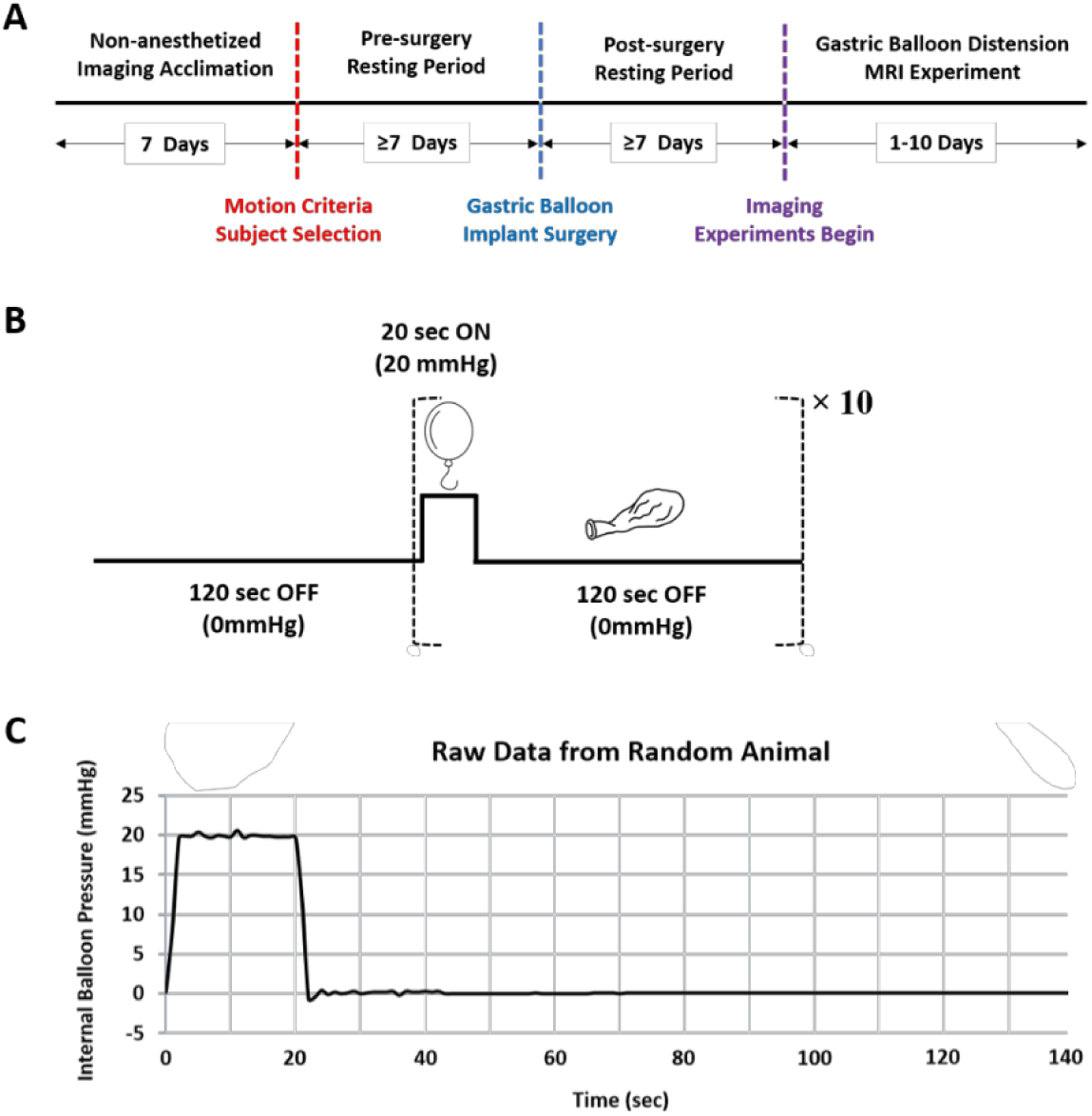
Relevant timelines / time courses used throughout the experiment. **(A)** Experimental timeline. The experiment can be roughly divided into three sections: 1) pre-surgery training, 2) gastric balloon implantation surgery, 3) gastric distension imaging. **(B)** Idealized time course of barostat schedule. Each gastric distention scan lasts for 1520 seconds: a starting 120-second-long non-pressurized state followed by a 140-second periodic schedule of 20 seconds at 20 mmHg and 120 seconds at 0 mmHg. The 140-second period repeats 10 times. **(C)** Representative time course of the 140-second ON-OFF pressure cycle. The attached graph depicts real-time barostat pressure readings (sampling rate 1 Hz) from an arbitrary animal during an imaging session. As can be seen, the pressure rise time and fall time are both ∼2 seconds.

### 2.4. BOLD fMRI – gastric balloon distension experiment

#### 2.4a. MRI parameters

MRI experiments were performed on a 7T Bruker 70/30 BioSpec running ParaVision 6.0.1 (Bruker, Billerica, MA) in the High Field MRI Facility at Pennsylvania State University. Radiofrequency (RF) transmission and signal reception were achieved with a volume quadrature coil (RAPID Biomedical) and 3-channel surface array coil (RAPID Biomedical), respectively. T2-weighted anatomical images were acquired using a rapid acquisition with relaxation enhancement (RARE) sequence with the following parameters: repetition time (TR) = 3.0 s, echo time (TE) = 40 ms, matrix size = 256 x 256, field of view (FOV) = 32 x 32 mm^2^, number of slices = 20, slice thickness = 1mm, averages = 6, RARE factor = 8, interleaved, anisotropic resolution = 125 x 125 x 1000 μm^3^. T2*-weighted functional images were acquired using a single shot GE-EPI sequence with the following parameters: TR = 1.0 s, TE = 15 ms, matrix size = 64 × 64, field of view = 32 × 32 mm^2^, number of slices= 20, flip angle = 60°, slice thickness = 1 mm, interleaved, anisotropic resolution = 500 x 500 x 1000 μm^3^. For acclimation-related imaging sessions (Day 5-7), functional data included four consecutive 600-volume scans per imaging session; for gastric distention imaging experiments (Methods 2.4c), functional data included two consecutive 1520-volume scans per imaging session. The spatially coincident anatomical and functional slice packages covered most of the brain: at least 2 mm and 3 mm of caudal olfactory bulb and rostral cerebellum, respectively, were included in the coverage. Note that all ‘consecutive’ functional scans acquired within the same imaging session are separated by approximately 5 minutes of preparatory MRI sequences (e.g. shimming).

#### 2.4b. Gastric balloon barostat

All gastric distension imaging experiments were preceded by ∼4 hours of fasting to minimize food content in the fundus. Prior to imaging, animals were first set up in a restrainer similar to the model used during the seven-day acclimation period. This imaging-specific restrainer included slight modifications to incorporate tubing that could integrate with the silicon catheter exiting at the rodent’s scapula (Figure 1C). After restrainer setup, animals were placed inside the MRI scanner, and tubing from the restrainer was interfaced with a balloon barostat (Distender Series IIR, G&J Electronics Inc.) to create a variable-volume, *closed system* filled with compressible air. As shown in Figure 1C, a plastic Y-connecter at the level of the implant’s catheter allowed for the barostat to measure and regulate internal air pressure separately. Briefly, a balloon barostat enables the experimenter to impose a fixed air pressure (relative to atmospheric pressure) acting against the interior wall of the balloon for a desired length of time. In order to first establish a positive pressure at the interior wall, an adjustable piston within the barostat is pushed towards the balloon to reduce the total volume of the balloon-tubing system. This decrease in total volume will cease once the pressure at the barostat’s measurement port (which has a constant cross-sectional area) reaches a user-specified value. When the volume of the balloon is finally at equilibrium, there will be a balance of forces acting on the balloon wall: elastic tension of the balloon wall + forces applied to external balloon wall = forces applied to internal balloon wall. In the gastric balloon context, external forces will be variable because the stomach wall can contract or relax around the balloon. Thus, after the balloon has reached volume equilibrium, increases or decreases in the applied external forces will result in the barostat’s piston being pulled away from or pushed towards the balloon, respectively, in order to maintain the desired pressure at the measurement port / balloon inner-wall. For the purposes of this project, the barostat was set to produce either a 0 mmHg (so-called “OFF” state) or 20 mmHg (so-called “ON” state) pressure. A diagram depicting key parts of a barostat is provided in the *Supplementary* (Supplementary Figure 1).

#### 2.4c. Gastric distension imaging

After setting up the barostat, the restrained animal was subjected to two 1520-second EPIs adhering to the following block design: 120 s OFF + (20 s ON + 120 s OFF) _x 10_, as depicted in Figure 2B. A user-programmed microcontroller (Arduino Uno) allowed the synchronization of barostat output with fMRI RF transmission (Figure 1D). Specifically, the microcontroller counted MRI-output TTL signals corresponding to the start of each new EPI volume and passed this data onto the barostat; different count numbers corresponded to different barostat-output pressure values. At the conclusion of the two fMRI scans, structural information was acquired using the aforementioned T2-RARE sequence. Across all animals included in this study, anywhere between one and five imaging sessions took place with each session producing a maximum number of 20 motion-passing ON-OFF cycles (10 cycles per scan, 2 scans per session). Supplementary Table 1 provides detailed information on the number of scans/epochs included for individual rats. For a given animal, imaging sessions were separated by at least 24 hours. After completing all imaging sessions, the animal was euthanized, and the abdominal cavity was opened to confirm that the balloon remained in its desired, gastric location. In all cases, real-time barostat volume curves informed the experimenter that leaks were not present in the balloon (Figure 2C). Additionally, ∼10 minutes prior to the 1^st^ scan of any given gastric distension imaging session, the balloon was transiently inflated using a sustained (45 seconds) barostat-driven pressure of 10 mmHg; after the 45 seconds, the pressure was returned to 0 mmHg. This pre-inflation was used to prime the balloon-barostat system for the initial periods of data collection.

### 2.5. Imaging data preprocessing

Imaging data preprocessing was implemented using in-house programs written in Matlab. Although previously described in greater detail ^23^, a brief description will be provided here. For a given functional scan (600 volumes or 1520 volumes), framewise displacement (FD) of every 20-slice volume was calculated to estimate scan-specific motion levels. Prior to generating FD measurements, the first 20 volumes of the scan were discarded to ensure that all volumes under subsequent consideration were acquired during MR signal steady state. In the context of non-anesthetized imaging acclimation, Day 7 was comprised of four separate 600 volume scans (580 following the aforementioned removal); animals that exhibited at least 550 (∼95%) volumes with an FD <0.2mm in three of the four scans were moved on to surgery.

In the context of the gastric distension imaging experiment, imaging sessions were comprised of two 1520 volume functional scans. At the outset of this preprocessing, a given functional scan had its 91^st^ through 1490^th^ volumes uniformly partitioned into 10 epochs; the first 90 volumes and final 30 volumes of a given functional scan were not used. Here, each *epoch* is defined as the 140-volume window of data starting 30 seconds before balloon distension and ending 90 seconds after balloon distension cessation (30+20+90). After labeling the start and end time of each epoch, all volumes were subjected to a stringent FD threshold of 0.1mm. Following FD calculations, all volumes with an FD > 0.1mm were tagged along with their immediate predecessor and successor volumes; double-tagged volumes were interpreted identically to single-tagged volumes. Following this FD-guided tagging procedure, only scans meeting the following criteria were included in further analysis: 1) no more than 20% (of 1520) of total volumes tagged, 2) no more than two epochs (out of the 10 total) contain tags coincident with the 20-volume distension period plus the final three and first three volumes of the pre and post-stimulus portions of the epoch, respectively (i.e. no tags in the 28^th^-53^rd^ volume of the 140 volume epoch). For each functional scan satisfying these criteria, an arbitrary non-tagged reference volume (belonging to the functional scan) was first manually aligned to a common rat brain anatomical template through rigid body transformation (between-modality alignment). All remaining non-tagged functional volumes were then subjected to automated rigid body motion correction and aligned to the original reference volume (within-modality alignment). FD-tagged volumes contained within the first 27 volumes (final 87 volumes) of the pre(post)-stimulus portion of each epoch were deleted and replaced with artificial data generated using a linear interpolation of the immediately bordering, non-tagged, motion-corrected and aligned volumes (Supplementary Figure 2). This approach allowed the 140-volume structure of the epochs to be preserved; for additional justification of this interpolation method, refer to Supplementary Figure 3. Subsequently, the resulting volumes were subjected to voxel-wise nuisance regression of motion parameters and within plane spatial smoothing (Gaussian kernel, FWHM = 1mm). Finally, each epoch’s 4D dataset was converted from arbitrary units (MRI output) to percent-change in signal. Specifically, for a given epoch, each voxel 𝑣_𝑖_ had its time series’ values 𝑣_𝑖𝑖_(𝑡) normalized to the temporal arithmetic mean μ_𝑖𝑖_ of its respective 30-volume pre-stimulation period: 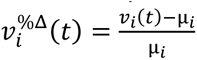. We define *pan-epoch* approaches as methods that do not differentiate between within-scan epoch numbering (e.g. 1^st^ epoch versus 2^nd^ epoch versus…versus 10^th^ epoch) and define *portion-epoch* approaches as methods that analyze specific subsets of within-scan epochs. Also, due to subtle differences in MRI slice acquisition from subject to subject, the two most anterior/rostral and posterior/caudal slices of fMRI data were not considered: only the interior 16 slices are used for analysis, and we refer to this total collection of 16 slices as *whole-brain*. Note that no white matter (WM) nor cerebrospinal fluid (CSF) nuisance regression was used in this analysis.

### 2.6. Imaging data post-processing

#### 2.6a. Two-level general linear modelling

After identifying gastric distension functional imaging scans satisfying motion-related criteria (Methods 2.5), voxel-wise two-level General Linear Models (GLM) were used to evaluate brain activation in response to balloon-induced distension of the rodents’ fundus. To prepare the data for the 1^st^ level analysis, all criteria-fulfilled scans were first broken down into their constitutive epochs (i.e. the fundamental object analyzed was a single 140-second epoch). An array composed consecutively of 30 zeros, 20 ones, and 90 zeros (representing the pre-stimulation, stimulation, and post-stimulation periods, respectively) was utilized to compute voxel-wise coefficients for each individual epoch (i.e. stimulation paradigm). This binary array was then convolved with a simulated hemodynamic response function (two linearly combined gamma functions with a resulting FWHM of 2.4s and time-to-peak of 2.9s). For a given epoch, the following model was used for each voxel of the brain:

*1^st^ – level analysis:*

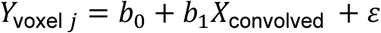

where the response variable 𝑌_voxel𝑗_ denotes the 1×140 fMRI time series for some voxel during the epoch, and X_convolved_ denotes the 1×140 regressing template time series (HRF convolved stimulation paradigm, Supplementary Figure 4). For each animal, this approach will generate multiple estimated 𝑏_1_ coefficients (one for each epoch) specific to a voxel. Supplementary Table 1 can be used to determine the number of voxel-specific 𝑏_1_ coefficients for each animal. After accumulating all voxel-specific and epoch-specific 𝑏_1_ coefficients associated with each animal, 2^nd^ level analysis involved voxel-wise linear mixed models: Matlab function ‘fitlme’. All coefficient estimates generated using this *Linear Mixed-Effect* (LME) function are subsequently referred to as *LME-derived*. To account for the repeated-measures (i.e. multiple voxel-specific 𝑏_1_ coefficients per subject), a *Subject* variable was incorporated and modeled as a random variable (random intercept): *Response* ∼ 1 + (1|*Subject*). Here, the modeled *Response* variable is comprised of all 𝑏_1_ coefficients previously generated from the 1^st^ level analysis. For clarity, this voxel-specific model can be conceptualized as:

*2^nd^ – level analysis:*

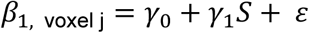

where 𝛽_1,voxel j_ is the 1×*m* list of all 𝑏_1_ coefficients (for a given voxel j) obtained from the 1^st^ level analysis, 𝛾_0_ is a fixed-effect average for all 𝑏_1_ coefficients of a specific voxel j, 𝛾_1_ is a 1×*n* array of estimated values for subject-specific intercepts, and 𝑆 is an *n*×*m* matrix of categorical indexing and zeros (with *m* representing the total number of epochs analyzed across all *n* unique animals). After performing the 2^nd^ level analysis, a univariate t-test was carried out on each voxel-specific 𝛾_0_ estimate. Specifically, we used the null hypothesis of 𝛾_0_ = 0 and recorded the resulting p-value in each voxel. To account for the issue of multiple comparisons, within-slice cluster-extent inference was performed using the following criteria: p-value < 0.005 and within-slice cluster size > 10. Under this framework, we claim that a *voxel-cluster* is significant at an 𝛼-level of 5%. The above methodology can be used with pan-epoch or portion-epoch approaches. To make claims about grey matter (GM) dominant Regions of Interest (ROIs), the whole brain was parcellated into 80 bilateral ROIs using the Waxholm Space atlas ^24^ for ROI-wise analysis (these GM-ROIs were distinct from defined WM and CSF masks). Importantly, the 80 defined bilateral ROIs and WM/CSF masks *did not* fully partition the brain: i.e. there were areas of the brain that were not assigned to GM, WM, or CSF classifications. The choice to leave these voxels as “Undefined” is motivated by the observation that known GM nuclei / WM tracts / CSF regions (provided by high resolution structural atlases) intersecting these volumes are typically smaller than the voxel size (500 x 500 x 1000 μm^3^) itself; accordingly, the name assignment scheme for these undefined voxels is objectively ambiguous. Note, although cluster inference only formally permits the claim that there is a 95% simultaneous confidence that every significant cluster contains at least one truly activated voxel, for linguistic convenience, we will refer to any GM-ROI that contains at least four voxels from any number of significant clusters as being an “*activated ROI*”. Similarly, any voxel that belongs to a significant cluster will be referred to as an ‘*activated voxel*’. Consequently, there can be activated voxels *not* contained by any GM-ROI or WM/CSF mask. To quantitatively describe these undefined voxels that are *also* activated, custom masks are used when appropriate (Results 3.1).

#### 2.6b. Spatial Independent Component Analysis

In a separate analysis, all 140-volume epochs across all scans and animals were used to identify brain activation patterns with a model-free method. This was achieved by group spatial independent component analysis (sICA) using the GIFT toolbox (Group ICA Of fMRI Toolbox) ^25^, where all epochs were pooled together. Importantly, sICA was applied to the 4D-dataset specific to the interior 16 slices (rather than all 20 slices). As with the GLM analysis, only epochs with sufficiently low motion during the stimulus and peri-stimulus window were included. Twenty maximally statistically independent spatial activation patterns were acquired using an infomax algorithm. This process was repeated times with random initial values to ensure the reproducibility of the resulting components. Of the twenty total spatial patterns, only patterns putatively of neuronal origin were selected for further analysis. For each of these spatial ICA components, spatial masks were generated using a z-value threshold ≥ 5. For a given spatial component, the resulting z-value threshold-mask was used to generate representative time courses. Specifically, for a given animal and a given epoch, the arithmetic mean was calculated from the time series of all voxels intersecting the threshold-mask to produce an epoch-specific 1x140 mean time series. This process was repeated across all animals and epochs. To calculate a representative ‘group average’ time series of a given spatial component, a linear mixed model (with a random intercept specific to each subject) was again used. In this context, though, each individual time point 𝑡 was separately modeled using the corresponding values from the epoch-specific mean time series. In general, then, one can view each value from the 1x140 ‘group average’ time series as being derived from the following model:

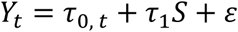

where 𝑌_𝑡_is the collection of all epoch-specific mean time series’ values at time 𝑡. Here, the 𝑡^th^ value of the representative ‘group average’ time series for a given ICA-generated spatial component is 𝑟_0, 𝑡_. 𝑟_1_ and 𝑆 adhere to the aforementioned (Methods 2.6a) structure. The above methodology can be used with pan-epoch or portion-epoch approaches.

#### 2.6 c. Accessory analyses of GLM output

Two accessory analyses were performed on the GLM data output. Recall, during the imaging procedure, each individual scan was comprised of 10 consecutive epochs. We therefore sought to determine if the brain response to stimulus varied across epoch number. To detect the presence of an epoch-dependent BOLD response, we first identified all activated ROIs from a pan-epoch GLM analysis. For each activated ROI, we made a corresponding spatial mask of any contained activated voxels: a so-called ‘activated voxel mask’. Note, then, that this analysis was restricted to only activated voxels intersecting with activated ROIs. Next, all epochs were separated on the basis of their chronological appearance during the imaging procedure: one pool of 1^st^ epochs, one pool of 2^nd^ epochs, etc. Then, for a given Category *i* of epoch number, the following procedure was performed:

- Choose any activated voxel mask 𝐾, which is comprised of 𝐽 activated voxels.
- Apply the mask to the corresponding 4D dataset of any epoch (across all scans and animals) belonging to the *i* ^th^ Category.
- For any such epoch, take the arithmetic mean 𝜇_𝑗_ of each within-mask activated voxel 𝑗’s 1x140 time series’ values from 𝑡 = 31 to 𝑡 = 55, and calculate the resulting arithmetic mean M_K_ across all such voxels: 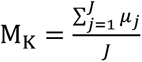. For clarification, 𝜇_𝑗𝑗_ is the arithmetic mean of the 25-second window coincident with the positive region of the HRF, as shown in Supplementary Figure 4.
- Repeat Step 3 across all animals and scans. This will result in a large pool of many M_K_ associated with the particular voxel mask 𝐾. A representative group average of M_K_ is then computed using a linear mixed model that is similar to the second-level GLM analysis. The resulting estimated coefficient is denoted by A_K_.
- Repeat steps 1-4 until the LME-derived group representative average A_K_ is generated for each activated voxel masks.
- Finally, the arithmetic mean of all of the different A_K_ (one for each mask) is calculated, along with the standard error of the mean.

Steps 1-6 can then be repeated for all epoch categories in order to generate an ‘*average*’ BOLD response of all activated ROIs identified by the pan-epoch GLM analysis: one for each category of epoch.

For the second accessory analysis, each activated ROI (as determined by any 2^nd^-level GLM output adhering to either a portion-epoch or pan-epoch approach) had a representative group average 1x140 time series calculated. These time series were generated using a similar approach to the first accessory analysis, but whole time series (rather than arithmetic means *of* time series) were treated as the primary objects of interest. Additionally, the statistical model that generated each activated ROI’s representative group average time series resembled the sICA linear mixed model (Methods 2.6b), where each time point (of the 140) was estimated with its own, separate model.

## 3. RESULTS

In order establish a translatable neuroimaging model of visceral manipulation, we used an implanted gastric balloon to perform a block-design sensory-evoked fMRI experiment in non-anesthetized rats. To detect regions of the brain that respond to 20 seconds of gastric distension (predominantly localized to the fundus), hypothesis-driven (GLM) and model-free (sICA) analyses were conducted. More specifically, GLM was used to explore the hypothesized temporal model of a 20-second square wave (convolved with an HRF) coincident with gastric distension; conversely, the sICA analysis made no assumptions regarding the temporal structure of the brain’s response to gastric perturbations. As will be seen, despite the different analytical approaches, GLM and sICA results exhibited considerable overlap among identified regions of BOLD activation in response to gastric distension.

### 3.1. GLM Results

The first GLM analysis incorporated all motion-passing epochs from all scans and subjects. Figure 3A provides a summary of this pan-epoch, 2^nd^-level GLM output for the whole-brain. As described in Methods 2.6a, the value of each brain voxel in Figure 3A represents the corresponding voxel-specific 𝛾𝛾_0_ estimate. Here, a larger (in magnitude) number is interpreted as denoting a voxel whose observed time series (across all scans/animals) was, on average, better modeled by the hypothesized temporal profile provided in Supplementary Figure 4. This initial analysis revealed a modest response in the cerebellum and ventral midbrain but only minimal changes throughout the remainder of the brain. In order to assess the BOLD response dependence on within-scan epoch number, we calculated the average, whole-brain response of activated ROIs as a function of within-scan epoch number (Methods 2.6c). As seen in Figure 3B (individual data points are shown in Supplementary Figure 5), within-scan epoch numbers 1, 2, and 6 yielded the highest average response across all epoch categories; possible mechanisms explaining this phenomenon are explored later in *Discussion*. A linear mixed model was used to confirm that the ROI-average percent change in BOLD signal was significantly higher among the 1^st^-2^nd^-6^th^ epoch subset than the 3^rd^-4^th^-5^th^-7^th^-8^th^-9^th^-10^th^ epoch subset (p=2.91×10^-13^, Supplementary Figure 6). Because of these differences in response, we next employed a portion-epoch approach by restricting our GLM analysis to epochs belonging to the 1^st^, 2^nd^, and 6^th^ categories. The resulting whole-brain GLM output can be seen in Figure 3C along with its corresponding cluster-thresholded map presented in Figure 3D. In addition to voxel-specific 𝛾_0_ coefficient estimates shown in Figures 3C and 3D, a corresponding p-value map is also provided in the *Supplementary* (Supplementary Figure 7). When restricting the GLM analysis to the 3^rd^-4^th^-5^th^-7^th^-8^th^-9^th^-10^th^ epoch subset, the spatial extent of significant responses was considerably smaller (Supplementary Figure 8). Restricting the analysis to the 1^st^-2^nd^-6^th^ epoch subset, however, substantially increased the number of activated voxels. Activated ROIs included the inferior colliculus, cerebellum, various hippocampal structures, and the ventral midbrain; a corresponding anatomical labeling of Figure 3D is provided in Supplementary Figure 9. A spatial summary of all activated voxels derived from the GLM analysis of the 1^st^-2^nd^-6^th^ epoch subset can be seen in Table 1; 3D renderings of this data can be found in Supplementary Figure 10. Several group average time series of activated voxels within activated ROIs (Methods 2.6c) are provided in Figure 4: inferior colliculus (Figure 4A), cerebellum (Figure 4B), retrohippocampal region (Figure 4C), and ventral midbrain (4D). Because of the large percentage of activated voxels that do not coincide with any pre-defined spatial location (Table 1), the time series of post-hoc, custom ROI maps were also included (Figure 4C and 4D). The time series displayed clear BOLD signal rise (and fall) near gastric distension onset (and offset). All group-averaged time series of activated voxels within *defined* ROIs can be found in the *Supplementary* (Supplementary Figures 11-14).

**Figure 3:**
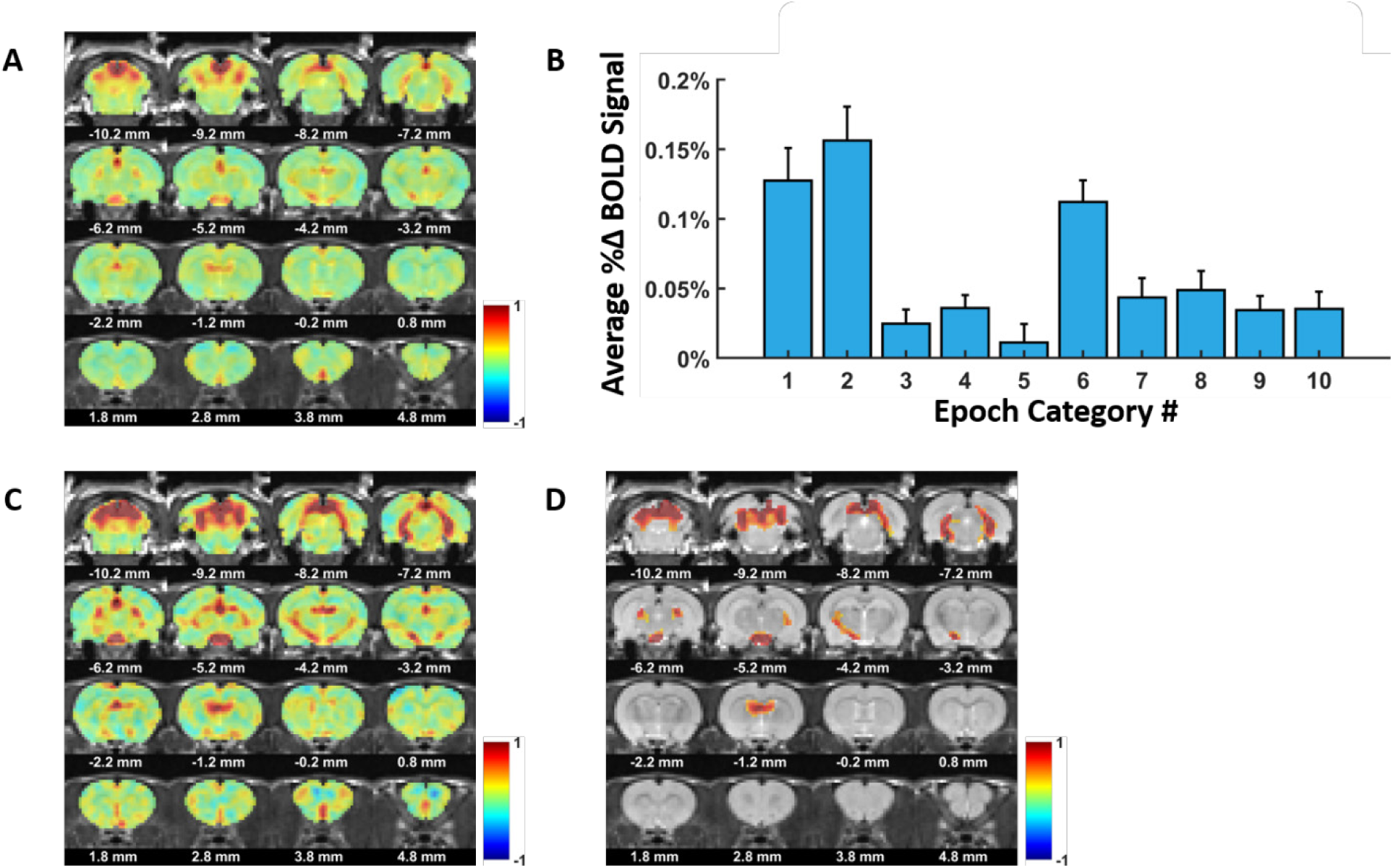
General linear modeling whole-brain output using pan-epoch and portion-epoch approaches. **(A)** Whole-brain representation of pan-epoch GLM analysis. Color intensity reflects the 𝛾𝛾_0_ estimate (second-level GLM) of each voxel. White numbers beneath each coronal cross section reflect the relative distance to bregma. **(B)** Bar plot capturing epoch-dependent BOLD responses. The height of each bar denotes a representative group average (Step 6 of Methods 2.6c) of all activated ROI responses associated with a particular epoch number; bars: standard error of mean. Stronger responses are observed during the 1^st^, 2^nd^, and 6^th^ epochs; weaker responses are observed during the 3^rd^, 4^th^, 5^th^, 7^th^, 8^th^, 9^th^, and 10^th^ epochs. To see the individual data points that were used in the LME models, refer to Supplementary Figure 6. **(C)** Whole-brain representation of 1^st^-2^nd^-6^th^ epoch subset GLM analysis. Restricting the GLM analysis to the high-response epochs results in a more widespread and intense brain activation. **(D)** Statistically-thresholded whole-brain representation of 1^st^-2^nd^-6^th^ epoch subset GLM analysis. This image depicts Panel C following cluster inference thresholding (Methods 2.6a). As indicated by the color bar, only positive 𝛾𝛾_0_ estimates survived in-plane cluster inference thresholding. The cerebellum, inferior colliculus, a variety of hippocampal structures, and ventral midbrain all show positive BOLD responses coincident with the 20-second 20-mmHg pressurization. For quantification of voxel-specific p-values (univariate hypothesis test of 𝛾𝛾_0_ = 0), refer to Supplementary Figure 7.

**Figure 4:**
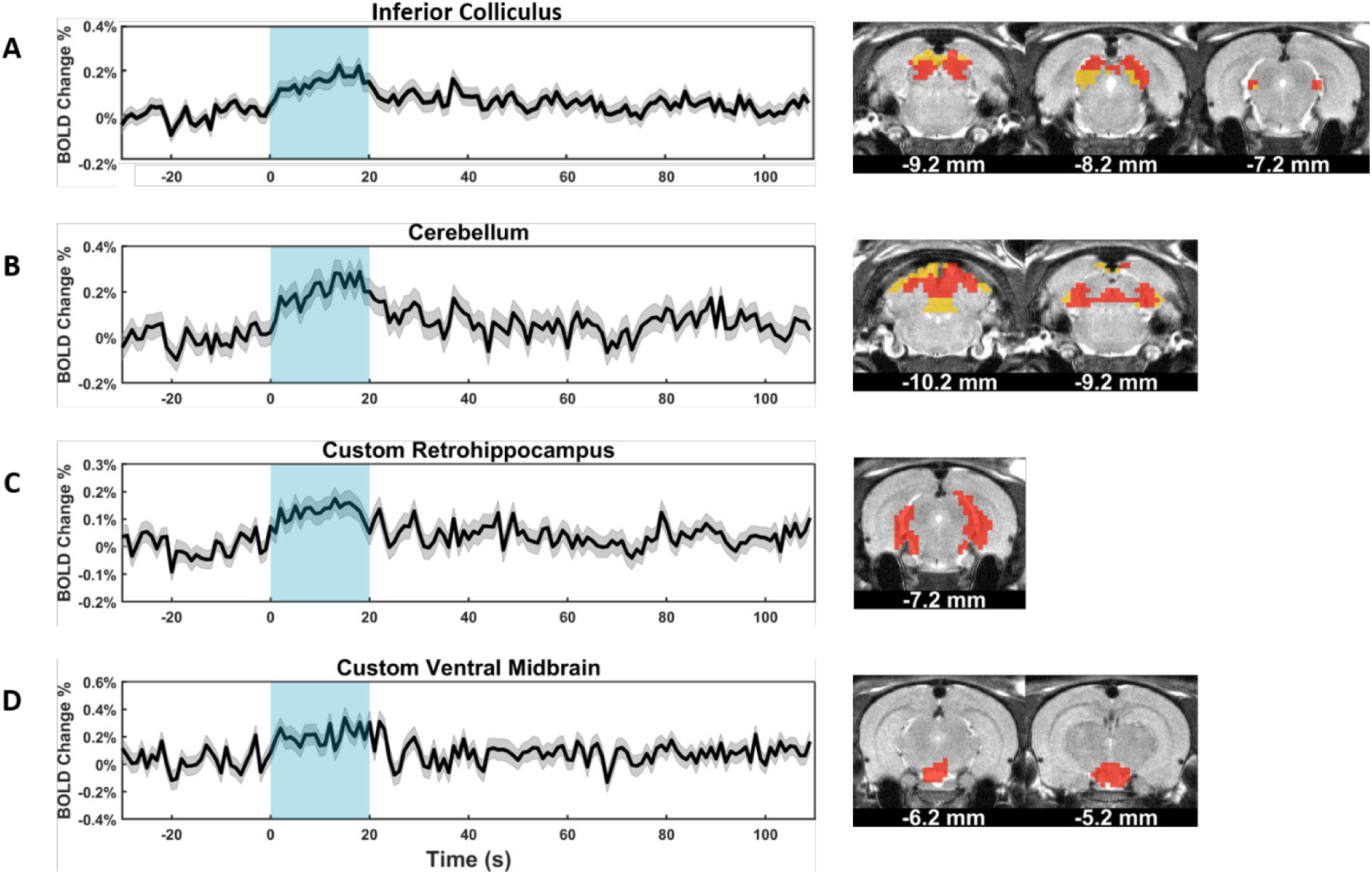
GLM-derived average time courses of selected regions. Panels A-D are all divided into a left and right column. The left column depicts LME-derived average time courses (black line) of select regions with widespread activation. The standard error of the model estimate associated with each time point’s LME-derived average is also included (gray shading). The blue box spanning from 0 to 20 seconds represents the 20-mmHg pressurization period. The right column depicts the corresponding ROI mask (yellow) and the activated voxels (red) contained within. The activated ROIs with the most pronounced spatial coverage were as follows: **(A)** inferior colliculus – 69% of mask occupied by activated voxels **(B)** cerebellum – 69% of mask occupied by activated voxels **(C)** custom retrohippocampus – by definition, 100% of mask is occupied by activated voxels **(D)** custom ventral midbrain – by definition, 100% of mask is occupied by activated voxels.

**Table 1:**
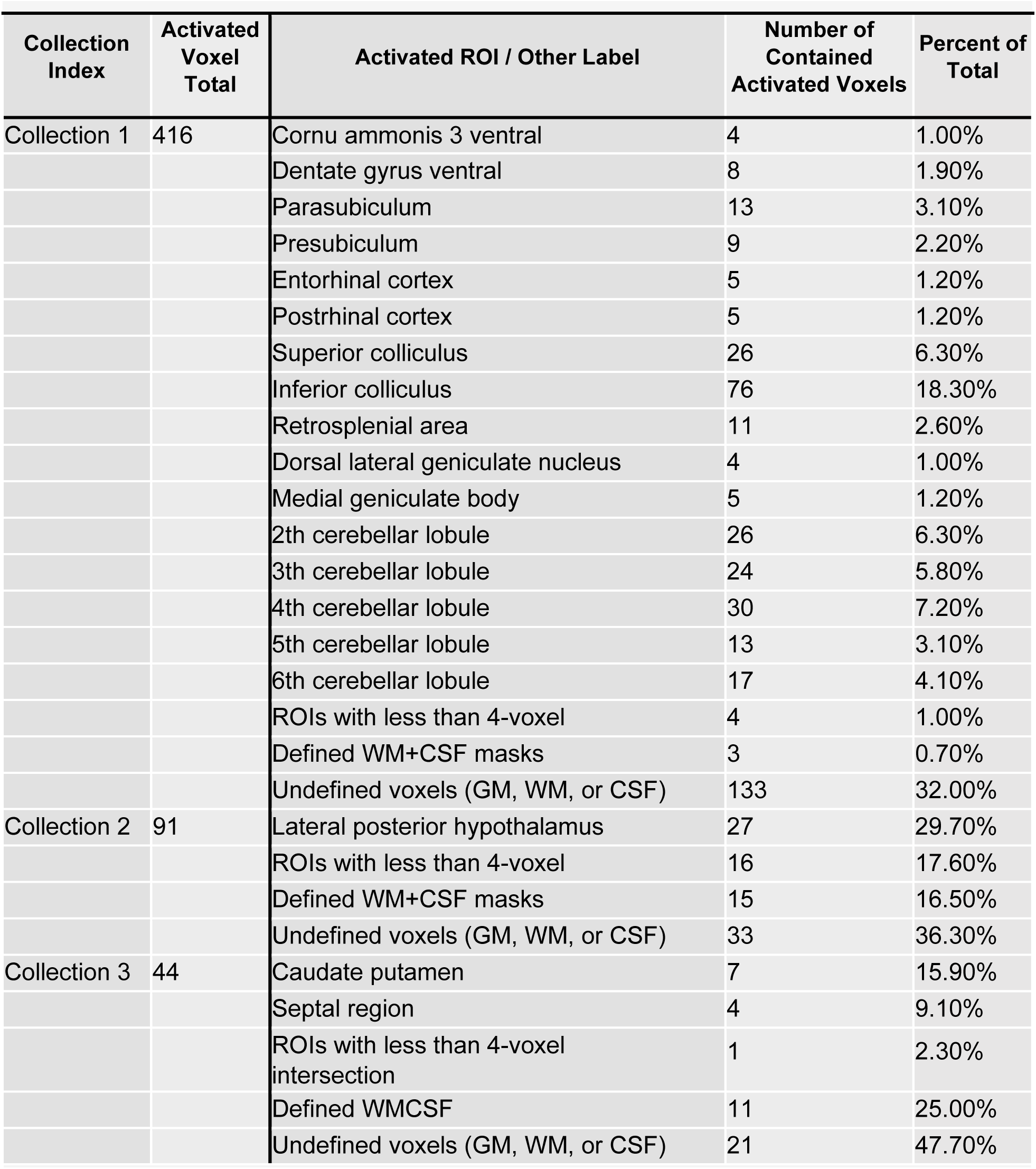
Summary of activated voxels from 1^st^-2^nd^-6^th^ epoch subset GLM analysis. This table provides an anatomical breakdown of the activated voxels output by the GLM analysis of the 1^st^-2^nd^-6^th^ epoch subset. Each collection (e.g. Collection 1) is comprised of 3D contiguous voxels that survived in-plane cluster inference thresholding. Two different renderings of these collections can **be found in Supplementary** Figure 9**. Note that not all activated voxels necessarily belong to a** defined GM ROI or WM/CSF mask (Methods 2.6a).

### 3.2. sICA Results

Using a two-level GLM analysis, we were able to assess how well a hypothesized hemodynamic response (Supplementary Figure 4) modeled the brain’s response to 20 seconds of gastric distension at 20mmHg. Importantly, with this approach, it is possible to *miss* brain responses whose temporal structures do not conform to the hypothesized model. To investigate the possibility of alternative brain responses, we performed whole-brain, group sICA, a method that presupposes no specific temporal structure. The results of Figure 3B again prompted the decision to only use sICA on data belonging to the 1^st^-2^nd^-6^th^ epoch subset. In order to identify voxels exhibiting temporal similarity, we used a large threshold value of z-value ≥ 5. Negative values (z-value ≤ -5) were not considered. 11 thresholded spatial maps, putatively of neuronal origin, were selected for further analysis. Representative group-averaged time series (Methods 2.6b) of four of the eleven maps are provided in Figures 5A-D. We see that the dentate gyrus (Figure 5A), midbrain (Figure 5B), cerebellum (Figures 5B & 5C), inferior colliculus (Figure 5D), and subicular complex (Figure 5D) all exhibit BOLD increases (peak amplitude between +0.1% and +0.2%) approximately coincident with gastric distension onset. The remaining 7 sICA components and accompanying time courses are provided in the *Supplementary* (Supplementary Figures 15-16).

**Figure 5:**
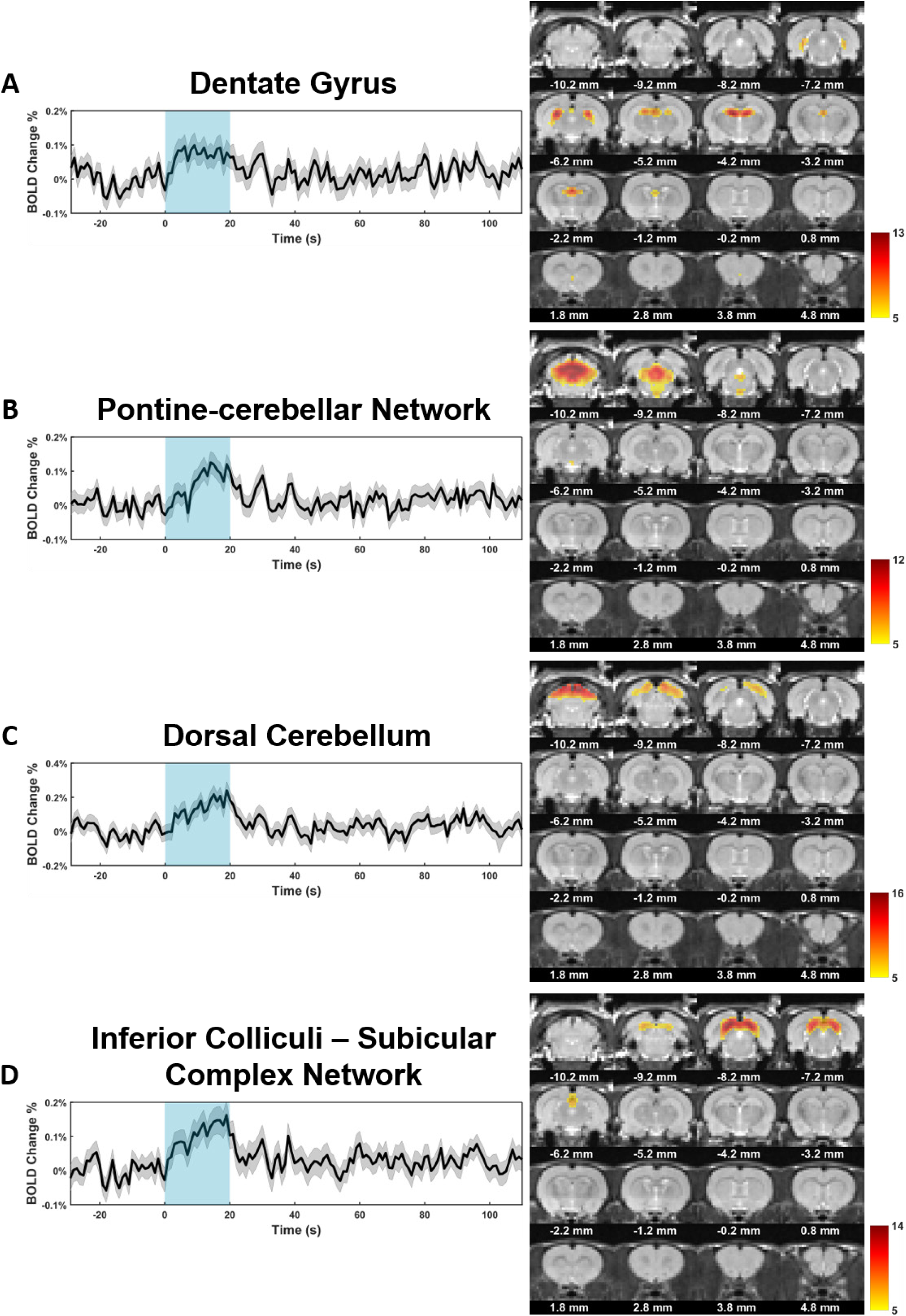
Representative sICA Components Derived From the 1^st^-2^nd^-6^th^ Epoch Subset. Panels A-D are all divided into a left and right column. The left column depicts LME-derived average time courses (black line), where each time point is modeled separately (Methods 2.6b), of spatial components calculated from the 1^st^-2^nd^-6^th^ epoch subset sICA analysis. The standard error of the model estimate associated with each time point’s LME-derived average is also included (gray shading). The blue box spanning from 0 to 20 seconds represents the 20-mmHg pressurization period. The right column depicts the spatial independent component maps from which the corresponding time courses were made. In accordance with the z-value thresholding criteria, all maps contain voxels with a positive z-value ≥ 5; the portions of the brain without yellow-red color intensities denote voxels whose z-values are < 5. The larger the z-value of two or more voxels, the greater the degree of temporal similarity among them. Each color bar is specific to its corresponding spatial map: the lower bound (yellow) is set to z-value = 5, and the upper bound (crimson) is set to the maximum voxel-specific z-value for that particular component. The sICA component maps are best categorized as follows: **(A)** dentate gyrus; **(B)** pontine-cerebellar network; **(C)** dorsal cerebellum; **(D)** inferior colliculi–subicular complex network.

## 4. DISCUSSION

This study sought to establish an unanesthetized animal model of visceral manipulation that concurrently captures global brain responses. BOLD fMRI permits the simultaneous, whole-brain measurement of deoxygenation states, which serve as proxies of neuronal activation. Therefore, using BOLD fMRI in combination with surgically implanted gastric balloons interfaced with a barostat, we imaged unanesthetized adult rats during periodic bouts of experimenter-controlled stomach distension. Initial analysis using general linear modeling revealed that the brain response to stomach distension exhibited an unexpected epoch dependency. Specifically, although maximal brain responses were observed at the beginning (1^st^ & 2^nd^ epochs) and middle (6^th^ epoch) of each scan, long intervals (> 400 seconds) of blunted responses followed both periods. Subsequent analysis (including sICA) ultimately focused *exclusively* on epochs that produced maximal brain responses; such refocusing enabled the identification of many more activated regions following gastric distension onset. We will first discuss the results under the interpretation that the high BOLD response of the 1^st^-2^nd^-6^th^ epoch subset is causally linked to underlying gray matter activation through classical neurovascular coupling. More specifically, we will treat high response regions of the brain as indicating sites of neuronal activation temporally correlated to gastric distension. Discussion thereafter will address possible confounds related to non-neuronal sources of BOLD signal change. Additionally, we will consider plausible mechanisms responsible for the observed epoch-dependent BOLD responses: high responses in the 1^st^-2^nd^-6^th^ epoch subset and low responses in the 3^rd^-4^th^-5^th^-7^th^-8^th^-9^th^-10^th^ epoch subset.

### 4.1. Implications of Activated Gray Matter Regions

The 1^st^-2^nd^-6^th^ epoch subset analysis revealed widespread BOLD activations in the cerebellum, inferior colliculus, ventral midbrain, and various hippocampal structures. These findings largely replicate previous balloon-based, gastric-distension PET and fMRI studies in humans ^5–13^ and anesthetized rats ^14,15^. Notably, though, several classically reported regions were absent such as the insula, cingulate gyrus, and the sensory cortices. However, many of these previously cited studies used paradigms that differed from ours in two relevant ways: duration of distension stimulus and extent of pain perception. More specifically, our 20-second distension period is quite short compared to other studies: ranging from 40 *seconds* ^10^ to as long as 60 *minutes* ^9^. Additionally, in these other studies, several balloon pressures/volumes used to distend the stomach were purposefully supra-pain threshold (induced by using greater pressures/greater volumes) in order to investigate a type of gastric pathology known as functional dyspepsia. Thus, our findings are probably specific to the *early-stages* of non-painful distension detection processes. In fact, on average, animal head motion frequency during gastric distension was only marginally (<1%) higher than during OFF periods (Supplementary Figure 3), which further suggests that our gastric manipulation did not induce pain.

Cerebellar and brainstem activation were observed in both the GLM and sICA analyses. Browning et al. (2014) reviews the cerebellum’s role in GI processing ^26^, noting that a variety of cerebellar projections indirectly (cerebellohypothalmic and cerebellovestibular) terminate throughout the brainstem, including the dorsal motor nucleus of the vagus (DMNV), a known stomach-modulating region ^26,27^. Our brain coverage was too rostral to capture the NTS and DMNV responses, but the sICA results (Figure 5) show a pontine-cerebellar network that clearly overlaps with the locations of the lateral (L) and medial (M) parabrachial nuclei (PBN). The MPBN and LPBN receive gustatory and viscerosensory information, respectively, from the NTS and relay this input to the lateral hypothalamus ^26^. Although the finding was relegated to the *Supplementary* (Supplementary Figure 12C), our distension paradigm revealed activation of the lateral and posterior portions of the hypothalamus, which partially coincided with our custom ventral midbrain mask (Figure 4D). Interestingly, the perifornical area of the hypothalamus is contained within these BOLD-activated hypothalamic areas; orexeginic (orexin-A) projections originating from this region terminate throughout the hippocampus onto gastric-distension-sensitive neurons that have been shown to regulate gastric motility ^28^. Finally, perhaps the most surprising result presented here is the overt activation of the inferior colliculus, a region classically associated with the auditory pathway ^29^. Spetter et al. (2014) has previously reported dorsal midbrain activation following gastric distension from water consumption (and chocolate milk), but no additional discussion of substructures (such as the inferior colliculus, which resides on the dorsal, inferior aspect of the midbrain) was provided ^30^. Nevertheless, Coleman et al. (1987) demonstrated that axons belonging to the dorsal column nuclei of the brainstem project to the inferior colliculus ^31^. Importantly, the dorsal column nuclei receive dense 1^st^-order projections from pseudounipolar spinal visceral afferents, a subset of which were recently shown to embed their opposite-end, organ-dwelling terminals within the muscle layers of the gastric fundus ^32^.

### 4.2. Addressing the possibility of non-neuronal sources of BOLD signal

In the normal biological setting, the BOLD fMRI signal strength is proportional to a water signal shaped by the relative presence of deoxygenated hemoglobin. Within a given voxel, the concentration of deoxygenated hemoglobin is tightly tied to ongoing hemodynamic parameters such as cerebral blood volume (CBV) and CBF ^33^. Therefore, although the BOLD signal provides an indirect measurement of neuronal activity, it is also sensitive to non-neuronal sources that affect deoxygenated hemoglobin state. Notable examples include certain anesthetic drugs like isoflurane ^34^ and hypertensive medications like sodium nitroprusside ^35^.

However, exogenous vasoactive molecules are not the only non-neuronal means of influencing the BOLD signal. In fact, even increases in intraabdominal pressure (IAP) can cause signal change. For example, using an inflatable pressure cuff around the waist of a mouse, pressurization onset was shown to be temporally coincident with brain motion occurring within the skull ^36^; such changes to brain position persisted until the cuff was depressurized. Importantly, brain motion can drive non-neuronal sources of BOLD signal variation through partial volume effects near GM-CSF and GM-vessel interfaces (leading to BOLD signal change in either direction). This finding is particularly relevant in the context of our stomach distension experiment because intragastric pressure (IGP) is known to have a strong linear correlation with intraabdominal pressure (IAP) ^37,38^. More specifically, our stomach distension experiment induces an intragastric pressure of 20 mmHg for 20 seconds, which necessarily brings about a proportional increase in IAP for the same window of time. In spite of this, it is doubtful that this factor dominantly contributes to the observed BOLD responses for the following reason.

Each of the 10 epochs within a given scan involve the same pressure application conditions. Consequently, if IAP increases (brought about by IGP increases) produced non-neuronal BOLD activations accounting for the majority share of signal increase, it seems reasonable to assume that this effect would be similarly replicated across every epoch. Clearly, this is not what we see, as the 1^st^-2^nd^-6^th^ epoch subset, on average, evokes a considerably stronger (and more widespread) response. It *is possible* that animals are, on average, elevating their IAP to greater levels on the first, second, and sixth epochs than the remaining epochs (thus enhancing the non-neuronal BOLD activation effect), but if IAP was elevated during the high response epochs, we could detect it with the barostat. This is done by assessing the position of the piston within the barostat; recall, the piston is pushed towards or pulled away from the balloon in order to clamp the internal balloon pressure at 20 mmHg. If the external pressure on the balloon increased (i.e. if the IAP rose), we would expect the piston to be further away from the balloon than if the external pressure was decreased. Using linear mixed modeling, we could not confirm the presence of such an effect: the piston position of the 1^st^-2^nd^-6^th^ epoch subset was not significantly different than the piston position of the 3^rd^-4^th^-5^th^-7^th^-8^th^-9^th^-10^th^ epoch subset (p = 0.1125, Supplementary Figure 17). Moreover, the tested coefficient had an estimated value of only 0.05518 mL (we converted piston distance change to barostat tubing volume change). A barostat volume - external pressure calibration curve was established by submerging a non-implanted gastric balloon (pressurized to 20 mmHg) at variable depths within a water-filled graduated cylinders and assessing how the barostat’s piston position changed. These curves revealed that a 0.05518 mL change in tubing volume equated to an IAP increase of only 1.72mmHg –3.34 mmHg. At these low-pressure differences, we find it unlikely that epoch-specific IAP increases are responsible for the epoch-dependent BOLD response. In summary, if IAP increases *were* the dominant source of BOLD activation, given that we have shown no significant IAP differences across epochs, we would be unable to explain the considerable signal contrast between high-response and low-response epochs. Therefore, we believe that the elevated responses of the 1^st^-2^nd^-6^th^ epoch-subset are dominantly driven by underlying gray matter activation.

### 4.3. Gastric accommodation as an explanation for epoch-dependent BOLD response

An alternative possibility that may explain the epoch-dependent BOLD responses involves the accommodation reflex of the stomach’s fundus. The accommodation reflex describes the phenomenon of reduced gastric muscle tone and increased wall compliance throughout the proximal stomach in response to inner-wall mechanical stimulation, which is observable in both rats ^39^ and humans ^40^. This reflex naturally arises in the context of food intake. For example, in the absence of any stomach volume changes, as an individual consumes food, IGP will rise. If the rate of consumption is sufficiently fast, IGP will continue to rise and, consequently, propel incompletely digested food from the stomach into the intestines. Fortunately, this premature release of stomach contents can be avoided by reducing smooth muscle cell tension in the stomach wall (i.e. reducing gastric tone), which allows for volume expansion while maintaining, or even lowering, IGP. Thus, the relaxation of smooth muscle ultimately *accommodates* the consumption of further food without sacrificing digestive efficacy. Interestingly, this reflex is predominantly achieved via a so-called *vagovagal pathway* whereby specialized neuronal afferent terminals (stretch receptors) embedded within the gastric wall respond to changes in muscle tension. These vagal afferents terminate in the brainstem (predominantly in the NTS) and initiate an efferent output originating from the DMNV ^27^. The preganglionic, parasympathetic projections from the DMNV terminate near neuronal nitric oxide synthase (nNOS) positive neurons that lay in close proximity to the fundus’ smooth muscle cells ^27,41,42^. Subsequent presynaptic release of acetylcholine from DMNV projections triggers the release of nitric oxide from the terminals of the nNOS+ cells onto the nearby smooth muscle, and relaxation ensues to complete the reflex.

To understand the relevance of this mechanism as it relates to the epoch-dependent BOLD response, recall that the gastric balloon is fabricated from non-compliant polyethylene. When the barostat is used to establish a 20-mmHg internal wall pressure on a *non-implanted* balloon, one can visually confirm that the balloon volume is essentially maximized: with the current fabrication approach, any further increases to internal pressure (i.e. > 20 mmHg) will result in virtually no additional balloon volume. Coupled with the ∼4-hour fasted state that animals were imaged under, one possible mechanism for the observed epoch-dependency can be outlined as follows. Just before the first distension onset of an imaging session’s first scan, the fundus exists in a contracted state (elevated gastric muscle tone) due to food absence. As the balloon expands into the fundus wall, the accommodation reflex initiates but only for a brief 20-second period before the 120-second OFF period begins. After pressure offset, the wall tension within the stomach wall has slightly reduced. Before the initial release of nitric oxide can be metabolized, though, the second distension onset begins. The balloon expands into the fundus wall that is now slightly less rigid due to the residual nitric oxide from the first epoch. Once again, an accommodation reflex is initiated, depositing even more nitric oxide into the fundus smooth muscle. At this point, the muscle tone is sufficiently relaxed so that the next three distension events (3^rd^, 4^th^, 5^th^ epochs) are unable to stretch the stomach wall any further *because the balloon has reached its maximal volume size.* Therefore, no additional reflexes are initiated. Once the sixth epoch has arrived, enough time has elapsed to remove the prior influence of nitric oxide on muscle relaxation, resulting in the return of enough muscle tone to induce an accommodation reflex during the 20-second pressure ON period. During the remaining four epochs (7^th^, 8^th^, 9^th^, 10^th^), the muscle tone is too weak and the balloon volume expansion too limited to induce an additional vagus-mediated stretch reflex. Due to intermediate MRI pulse sequences separating within-session EPI scans, when the first distension onset of the *second* within-session scan begins, enough time will have elapsed to return the fundus’ muscle tone to a sufficiently rigid state, and the argument repeats. Under this speculative framework, the large amplitude BOLD responses observed in the 1^st^-2^nd^-6^th^ epoch subset are due to stretch information being conveyed at the level of the stomach up to the brainstem and then propagating further rostrally to the cerebellum, midbrain, and subcortical structures. In contrast, the blunted BOLD responses observed in the 3^rd^-4^th^-5^th^-7^th^-8^th^-9^th^-10^th^ epoch subset are due to a *weak* stretching stimulus, leading to a proportionately weak brain response.

## 5. CONCLUSION

In the current paper, we demonstrated that periodic barostat-induced gastric distension combined with simultaneous BOLD fMRI can provide a neuroimaging model of visceral manipulation in non-anesthetized rats. Accompanying GLM and sICA analyses revealed widespread BOLD activation across the inferior colliculus, cerebellum, ventral midbrain, and a variety of hippocampal-related structures. The inferior colliculus finding is noteworthy, as there is minimal preexisting literature linking this structure to the broad category of interception, let alone to the stomach. This structure’s participation in processes extending beyond its classically associated role in the auditory pathway emphasizes the continued need for neuroscience to embrace the framework of multisensory functionality within brain nuclei. Most importantly, to the best of our knowledge, this experiment showed for first time ever that anesthetic agents are unnecessary for the neuroimaging of animals during experimenter-controlled perturbations of major organs. Not only does this dispel the concerns of anesthesia-induced signal confounds and associated translatability issues, but the acquired dataset clearly indicated an absence of motion-induced artifacts that should assuage often-cited worries of non-sedative approaches. The success of this novel experiment widens the space of possible paradigms focused on enriching the current understanding of brain-viscera interactions. For example, animals as subjects greatly expands the range of neuroscientific tools at one’s disposal. A clear direction to pursue is the direct placement of MRI-compatible recording electrodes within organ walls to access more refined measurements of organ behavior, which can better inform interpretations of the resulting brain responses. Further, the non-anesthetized state of the animal permits more biologically relevant investigations by enabling researchers to explore context-dependent interoceptive responses; such responses may otherwise be offline during anesthesia-induced unconsciousness. Ultimately, non-anesthetized animal imaging presents itself as a powerful approach to track and correlate whole-brain responses to ongoing visceral activity.

## Acknowledgments

We thank Drs. Shiying Li, Jiande Chen and Zhongming Liu for their technical support. The present study was partially supported by National Center for Complementary and Integrative Health (R01AT011665). The content is solely the responsibility of the authors and does not necessarily represent the official views of the National Institutes of Health.

## Author Contributions

SC and NZ designed the experiment. SC conducted all experiments. XH conducted primary analysis. DC conducted secondary analysis. SC designed primary equipment. TN designed secondary equipment. SC wrote the paper. SC, XH, and NZ edited the paper. NZ acquired the funding.

