## Supplemental Document for "Neuroimaging model of visceral manipulation in awake rat"

**Affiliations**:
1 The Neuroscience Graduate Program, The Huck Institutes of the Life Sciences, The Pennsylvania State University, University Park, USA
2 Department of Biomedical Engineering, The Pennsylvania State University, University Park, USA
3 Center for Neural Engineering, The Pennsylvania State University, University Park, USA
4 Center for Neurotechnology in Mental Health Research, The Pennsylvania State University, University Park, USA

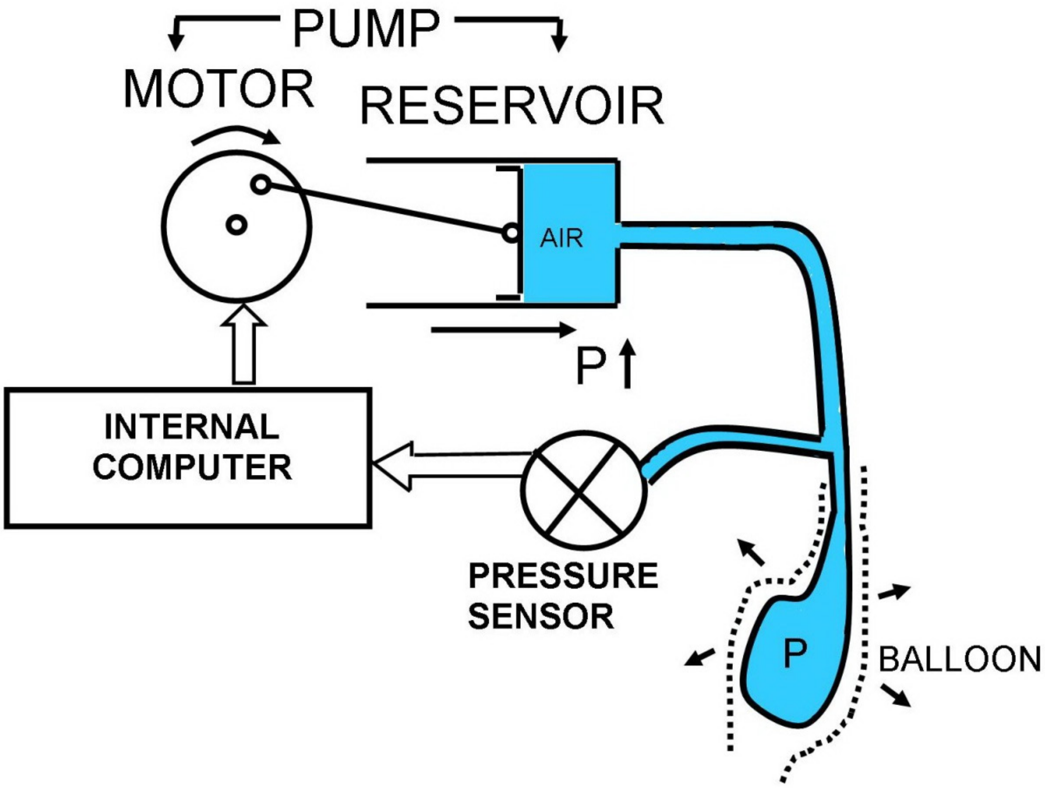


**Supplementary Figure 1: Schematic of barostat components**. The gastric implant's catheter is connected to two tubes via a Y-connector. These two tubes interface with two different barostat ports. One port houses a pressure sensor, and the other port houses a sealed air pump chamber, which can be likened to a mechanized syringe. Software tracks real-time pressure sensor data to determine whether or not the pump should be pushed further towards or pulled away from the balloon (in order to maintain the programmed pressure). In this experiment, the barostat was required to occupy two different states of sustained pressure throughout each 1520-second functional scan: 0 mmHg (LOW/OFF) and 20 mmHg (HIGH/ON).

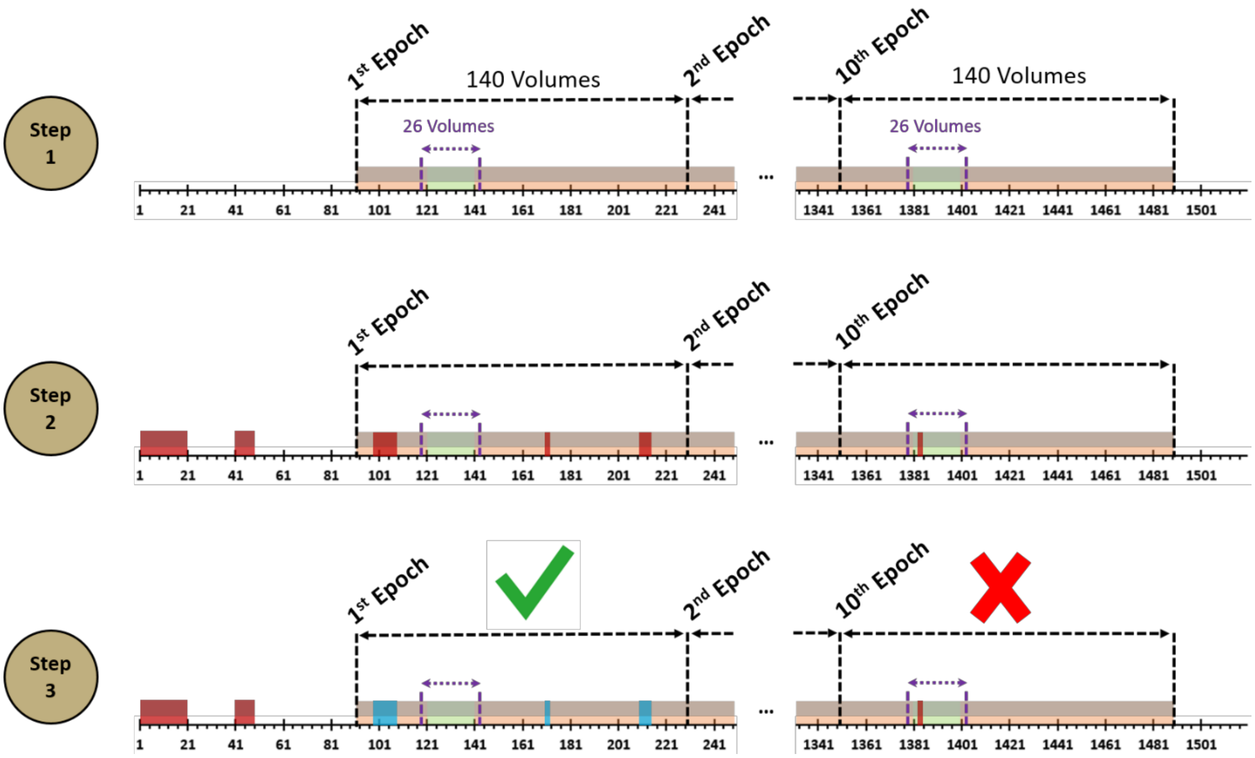


**Supplementary Figure 2: Description of gastric distension functional scans' data structure.** (*Step 1*) The first step in the processing of a gastric distension functional scan (1520 total volumes) involves partitioning the $91^{st}$-$1490^{th}$ volumes into ten distinct 140-volume epochs. In this illustration, the beige rectangles within each epoch (black dashed lines) represent periods of 0-mmHg pressurization; the accompanying green box represents the 20-mmHg pressurization state. Note that the $1^{st}$-$90^{th}$ and $1491^{st}$-$1520^{th}$ volumes are also coincident with a 0-mmHg pressurization state but are *not* involved in the epoch-partitioning process and, therefore, are not involved in the overall analysis. (*Step 2*) The entire 1520-volume scan is subjected to framewise displacement (FD) calculations. Volumes with FD-value > 0.10 mm are tagged (red box). Regions (within-epoch or outside-of-epoch) with FD-value > 0.10 are all used to determine if a scan has at least 80% of volumes below the motion threshold. (*Step 3*) Within each epoch, the 26-volume (purple arrow) "No Motion" region denotes periods of time that cannot have any FD value > 0.10 mm. If tagged volumes are inside an epoch and *not* within the No Motion region, interpolation (with the closest bordering non-tagged volumes used to determine the boundary conditions) on a voxel-to-voxel basis is used to replace the tagged volumes (blue box). If an epoch contains no above-threshold volumes in its No Motion region (e.g. $1^{st}$ Epoch), the epoch passes. If an epoch contains FD-tagged volumes in the No Motion region (e.g. $10^{th}$ Epoch), the epoch does not pass and is discarded. In order for a scan's epochs to participate in subsequent analysis, the given scan must have at least 8 out of 10 epochs pass. If 7 or less of the scan's epochs pass, then *none* of the epochs from this scan move on to further analysis.

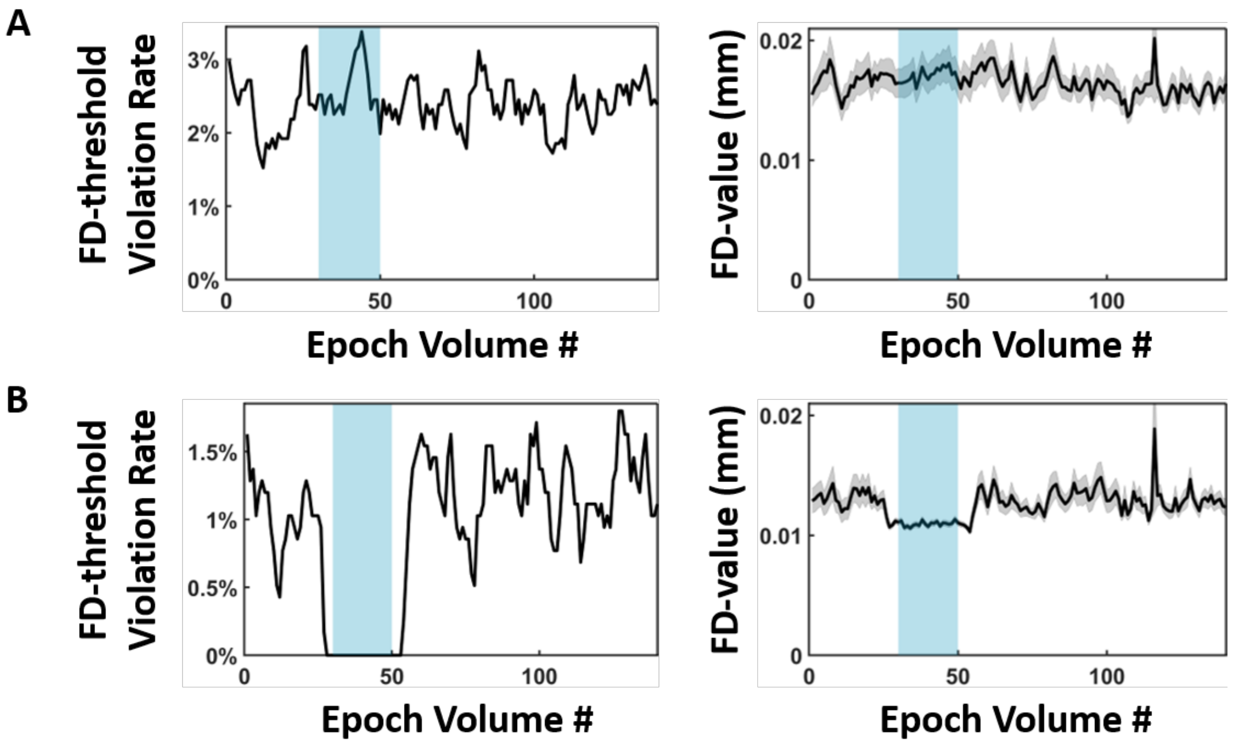


**Supplementary Figure 3: Using motion data to justify data interpolation**. **(A)** Graphs summarizing fMRI motion data across the collective pool of passing and non-passing epochs. (*Left*) 20 mmHg gastric balloon pressurization is denoted by the blue rectangle, which spans 20 volumes (equivalently spanning 20 seconds due to TR=1.0 second). The y-axis describes the percentage of all (passing/good and non-passing/not good) epochs that were above the FD-value threshold of 0.10 mm for a given volume. (*Right*) The arithmetic mean of each volume's FD-values across all (passing/good and non-passing/not good) epochs was plotted (black line); similarly, the corresponding standard error of the mean is illustrated for each volume (gray shading). **(B)** Graphs summarizing fMRI motion data across exclusively passing epoch. (*Left*) The y-axis describes the percentage of all passing epochs that were above the FD-value threshold of 0.10 mm for a given volume. By definition, 0% of passing scans have FD-threshold violations (FD-value > 0.10 mm) during the 28$^{th}$-53$^{rd}$ volumes. Across all other volumes (1$^{st}$-27$^{th}$ and 54$^{th}$-140$^{th}$), the FD-threshold violation rate is never larger than 2.0%. Further, the rate is relatively uniform in this volume range, suggesting that animal motion events are not biased towards any particular time. (*Right*) The arithmetic mean of each volume's FD-values across all passing epochs was plotted (black line); similarly, the corresponding standard error of the mean is illustrated for each volume (gray shading). As a consequence of the selection criteria, the average FD-value of the 28$^{th}$-53$^{rd}$ volumes are lower than the 1$^{st}$-27$^{th}$ and 54$^{th}$-140$^{th}$ volumes (~0.01 mm). (*Both*) Panel B data suggests that, across all passing epochs, motion is relatively uniform, and, in particular, supra-threshold motion is consistently infrequent in the 1$^{st}$-27$^{th}$ and 54$^{th}$-140$^{th}$ volumes. As such, it is unlikely that interpolation within the 1$^{st}$-27$^{th}$ and 54$^{th}$-140$^{th}$ volumes added any structured, artificial signal into the BOLD time series.

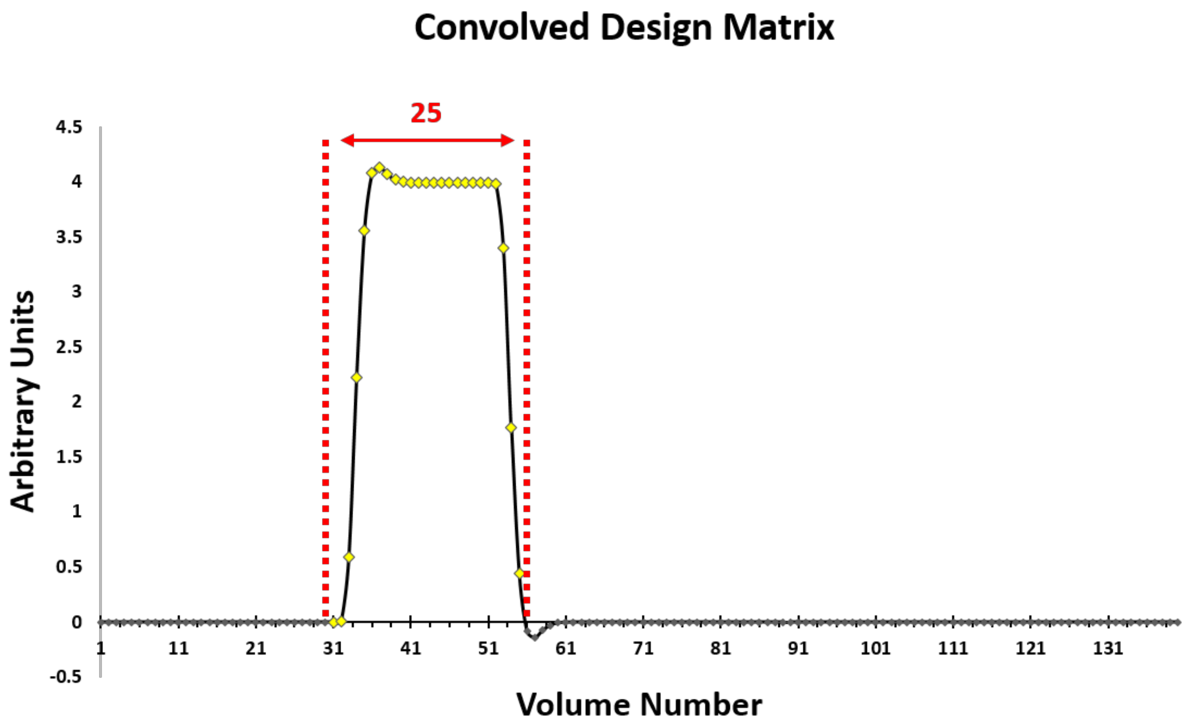


**Supplementary Figure 4: Visualization of the HRF curve used in the GLM analyses.** This figure depicts the 1x140 HRF time series used in the GLM analyses. It is generated by convolving a square wave (30 zeroes → 20 ones → 90 zeroes) with the linear summation of two gamma function. As can be seen, the resulting curve contains 25 positive values (denoted as yellow diamonds contained within the red, dashed lines) that reflect the hypothesized BOLD response to a 20-volume-long square stimulus function. The onset of this positive response begins at the 31$^{st}$ volume and persists until the 55$^{th}$ volume. This justifies section 2.6c's calculation of the arithmetic mean of the BOLD response over this window: the 31$^{st}$-55$^{th}$ volumes represent the period of positive response within the hypothesized model.

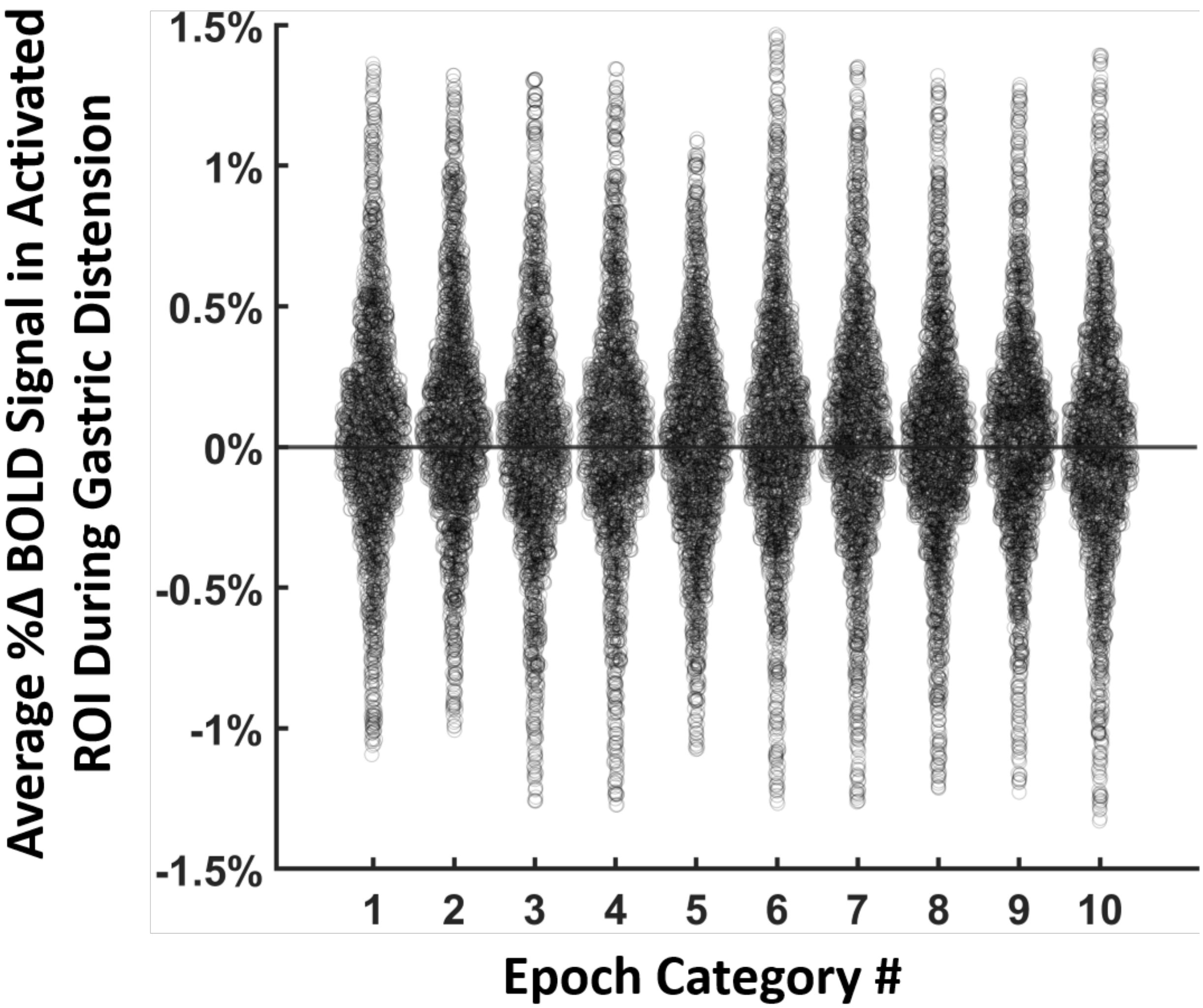


**Supplementary Figure 5: Individual data used to generate the bar plots of Figure 3B.** This figure depicts most of the data used to generate the bar plots in Figure 3B. For display purposes, for each epoch category, only 95% of the data closest to the mean (Step 6 of Methods 2.6c) is shown. For a given epoch category $i$, each data point represents the arithmetic average of the BOLD time-series of activated voxels within an activated ROI (determined by the pan-epoch GLM analysis) over a 25-second window (31st volume – 55th volume) for one animal, for one scan. Each data point for a given epoch is therefore equivalent to one instance of an $M_K$ (Step 3 of Methods 2.6c).

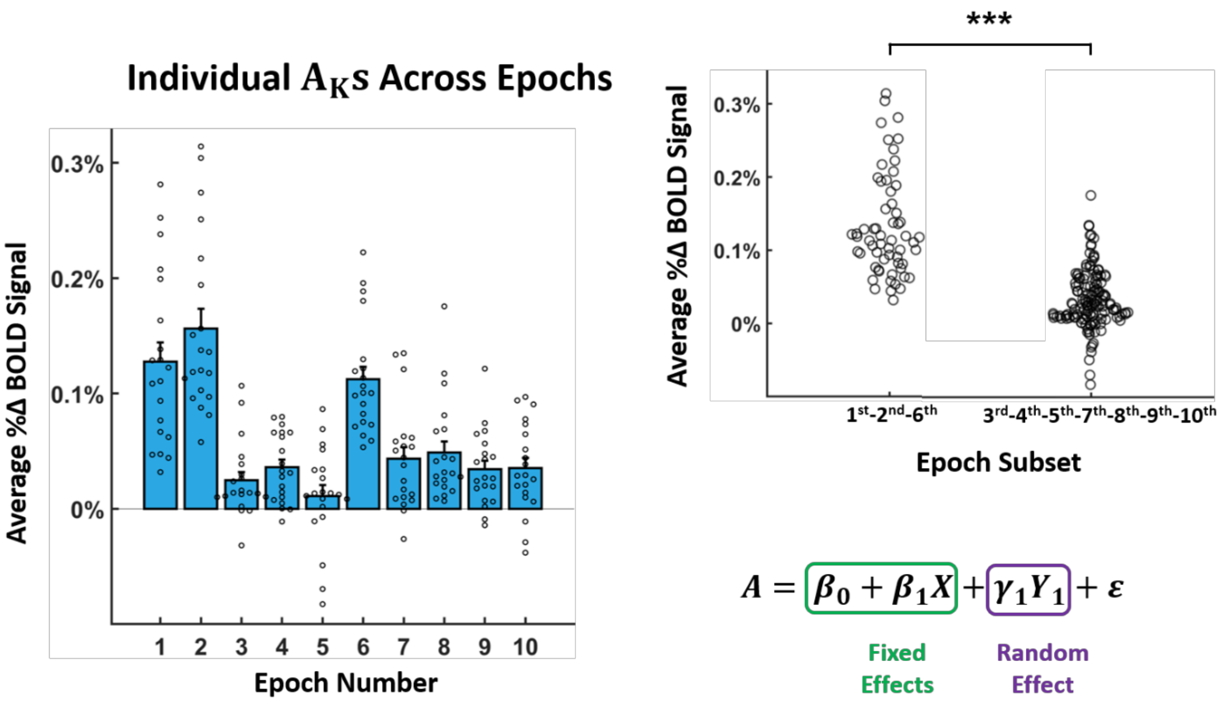


**Supplementary Figure 6: 1st-2nd-6th epoch subset has significantly larger average activated ROI BOLD response during gastric distension compared to 3rd-4th-5th-7th-8th-9th-10th epoch subset.** (*Left*) This bar plot is a replica of Figure 3B but with the addition of all individual LME-derived $A_K$s (Step 6 of Method 2.6c). The pan-epoch GLM analysis identified 20 activated ROIs. Therefore, for epoch number $i$, there are 20 different plotted values: $A_1^i, A_2^i, \cdots, A_{20}^i$. (*Top Right*) A collapsed version of the left bar plot is provided to emphasize the differences between the two epoch subsets. The 1st-2nd-6th epoch subset contains 60 $A_K$s ($A_1^1, \cdots, A_{20}^1, A_1^2, \cdots, A_{20}^2, A_1^3, \cdots, A_{20}^3$), and the 3rd-4th-5th-7th-8th-9th-10th epoch subset contains 140 similarly indexed $A_K$s. A linear mixed model (described next) reported that, on average, the activated ROI BOLD responses arising during the 1st, 2nd, and 6th epochs were significantly larger than the responses from the 3rd, 4th, 5th, 7th, 8th, 9th, and 10th epochs (p-value < 0.001). (*Bottom Right*) The general form of the linear mixed model used to test for the presence of activated ROI BOLD response differences between epoch subsets is shown. Two fixed effects and one random effect were used to assess whether or not the average activated ROI BOLD response during gastric distension of the 1st-2nd-6th epoch subset was significantly larger than the responses of the 3rd-4th-5th-7th-8th-9th-10th epoch subset. Fixed effects included a global average $\beta_0$ and an epoch-subset-specific coefficient $\beta_1$, where $X$ was a categorical variable taking on the value of either 0 (if $A_K^i$ was from the $i = 1, 2, 6$ epochs) or 1 (if $A_K^i$ was from the $i = 3, 4, 5, 7, 8, 9, 10$ epochs). The random effect encoded a random intercept for each activated ROI. The model is thus structured to treat each activated ROI $K$ as being sampled 10 different times: once for each epoch. We rejected the null hypothesis of $\beta_1 = 0$ ($p=2.91\times10^{-13}$).

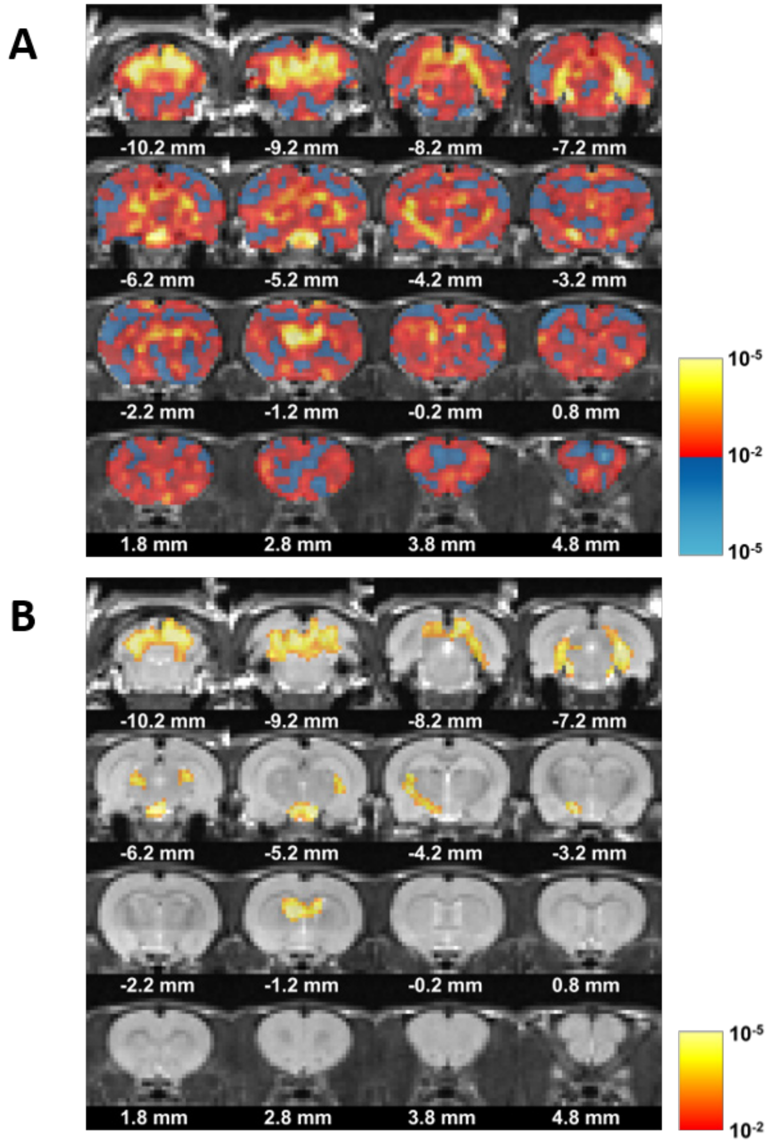


**Supplementary Figure 7: Voxel-specific p-value representation of 1st-2nd-6th epoch subset GLM analysis. (A)** Whole-brain depiction of univariate hypothesis test results. This panel depicts the p-value output across all voxels for the univariate hypothesis test of $\gamma_0 = 0$ (Methods 2.6a). The color bar indicates two types of voxels: those with $\gamma_0 < 0$ and those with $\gamma_0 > 0$. If $\gamma_0 > 0$, then the red-to-yellow gradient color scheme is applied; if $\gamma_0 < 0$, then the dark-blue-to-light-blue gradient color scheme is applied. The white digits beneath each coronal brain cross section reflects the location of the slice relative to bregma. **(B)** P-value representation of voxels surviving cluster thresholding. This image replicates Figure 3D, but the color bar intensity here represents p-values instead of model coefficient estimates. Note that no voxel from Panel A with $\gamma_0 < 0$ survives within-slice cluster inference thresholding.

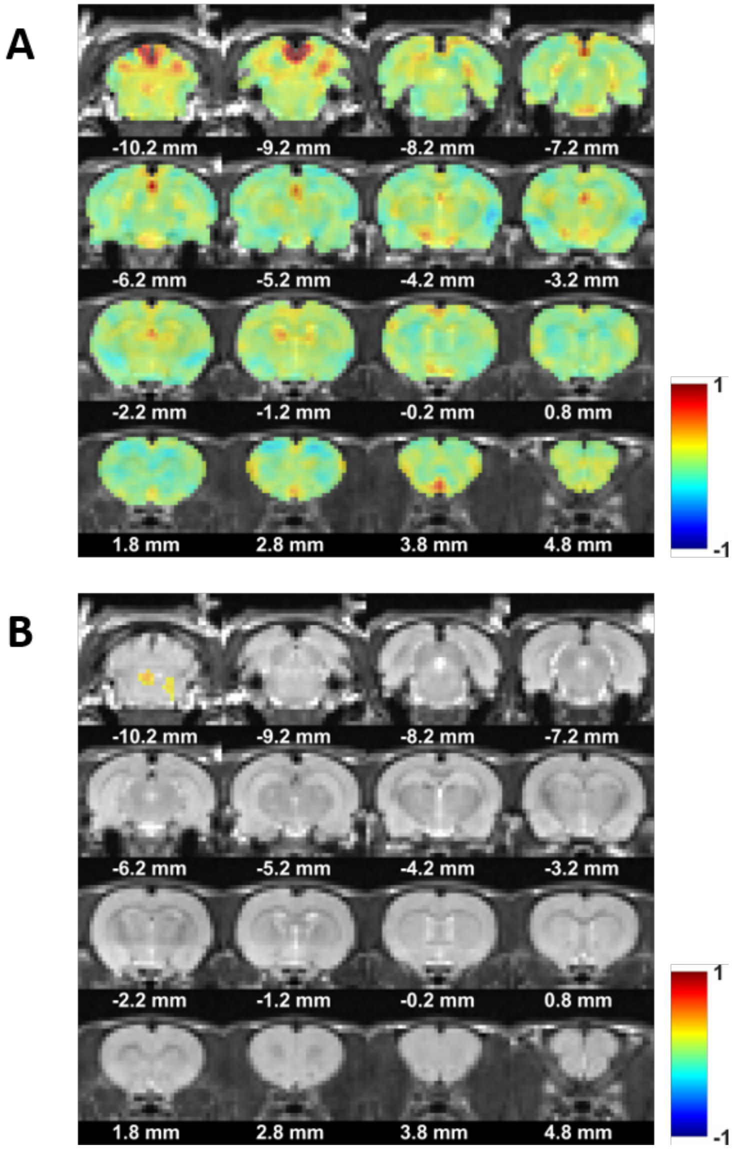


**Supplementary Figure 8: GLM analysis of 3$^{rd}$-4$^{th}$-5$^{th}$-7$^{th}$-8$^{th}$-9$^{th}$-10$^{th}$ epoch subset. (A)** Whole-brain depiction of voxel-specific $\gamma_0$ estimates for 3$^{rd}$-4$^{th}$-5$^{th}$-7$^{th}$-8$^{th}$-9$^{th}$-10$^{th}$ epoch subset. This panel represents the non-thresholded output of the portion-epoch GLM analysis of the 3$^{rd}$-4$^{th}$-5$^{th}$-7$^{th}$-8$^{th}$-9$^{th}$-10$^{th}$ epoch subset. Color intensity is used to signify the $\gamma_0$ value output by the second-level GLM analysis for each voxel. The white digits beneath each coronal brain cross section reflects the location of the slice relative to bregma. **(B)** $\gamma_0$ estimates of voxels surviving cluster thresholding. This figure identifies which areas of the brain survived within-slice cluster inference thresholding (Methods 2.6a). Compared to Figure 3D (Results 3.1), there is considerably less activation of the brain in response to gastric distension during the 3$^{rd}$, 4$^{th}$, 5$^{th}$, 7$^{th}$, 8$^{th}$, 9$^{th}$, and 10$^{th}$ epochs. Nonetheless, marginal activation is observed in the posterior of the brain, near the ventral cerebellum and dorsolateral brain stem.

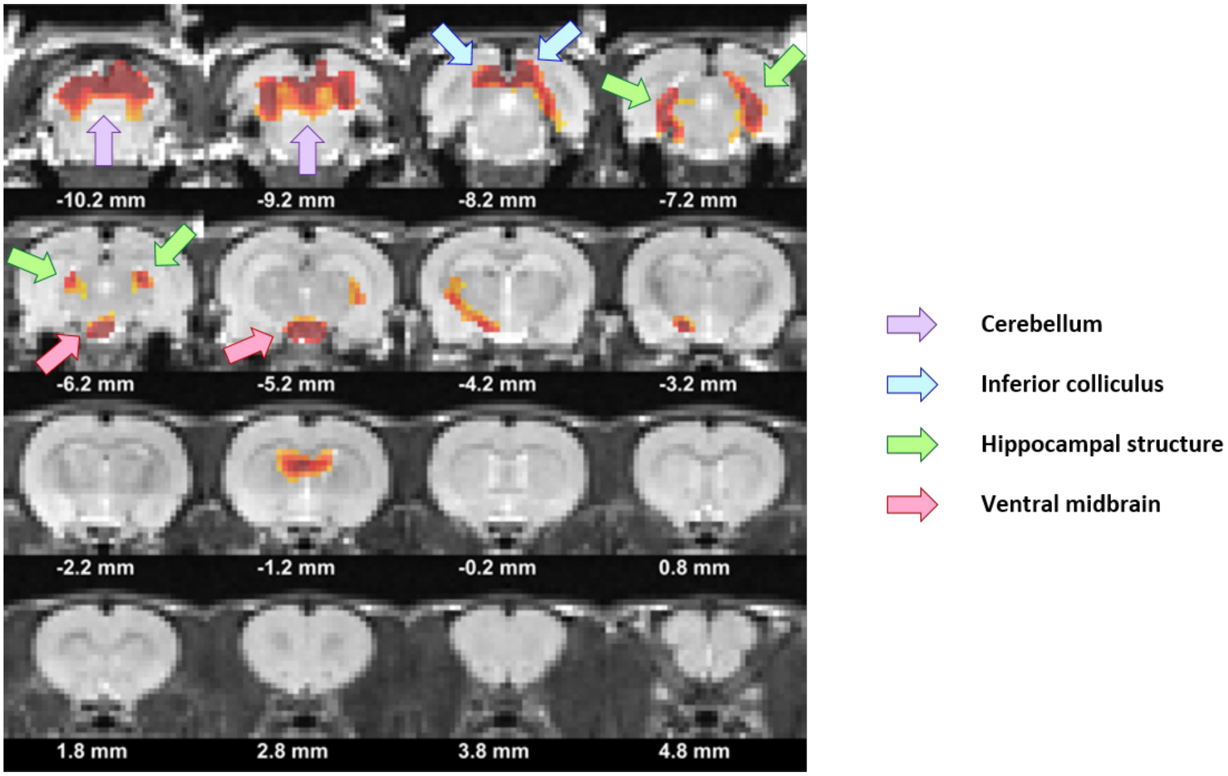


**Supplementary Figure 9: Anatomical Annotations of Cluster-Thresholded $1^{st}$-$2^{nd}$-$6^{th}$ Epoch Subset GLM Analysis.** This figure is a replica of Figure 3D (Results 3.1) but with additional anatomical annotations: cerebellar activation is indicated by purple arrows; inferior colliculi activation is indicated by blue arrows; a variety of hippocampal activations are indicated by green arrows; and ventral midbrain activation is indicated by red arrows.

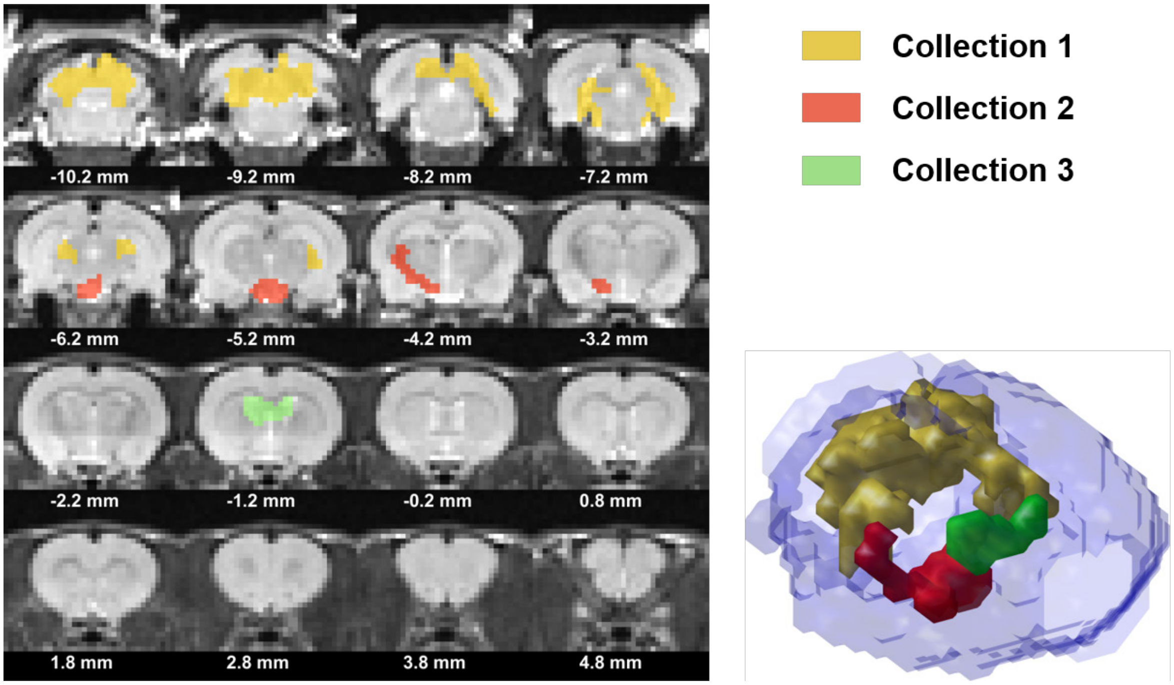


**Supplementary Figure 10: Identifying 3D-contiguous collections of thresholded clusters for 1st-2nd-6th epoch subset GLM analysis.** (*Left and Top Right*) This figure shows that there are three distinct 3D-contiguous collections of clusters following in-plane cluster inference thresholding. Along the caudal-rostral axis, we have Collection 1 (yellow), comprised predominantly of the cerebellum, inferior colliculi, and hippocampal regions, Collection 2 (red), comprised predominantly of the ventral midbrain, lateral and posterior portions of the hypothalamus, and white matter (internal capsule and cerebral peduncle), and Collection 3 (green), comprised predominantly of CSF fluid (third ventricle and lateral ventricle) and hippocampal white matter (such as the fimbria). (*Bottom Right*) A glass-brain was used to visualize the spatial relationships amongst the three collections in 3D.

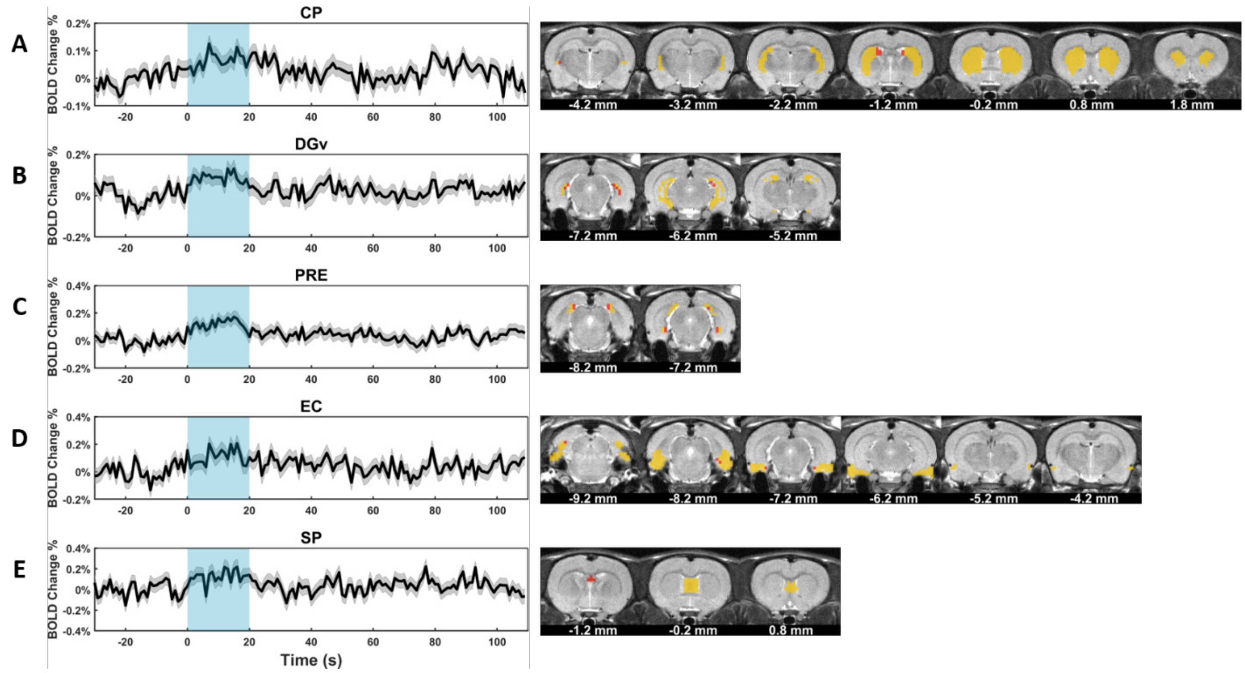


**Supplementary Figure 11: Additional GLM-derived average time courses for all active ROIs – Part I.** Panels A-E are all divided into a left and right column. The left column depicts LME-derived average time courses (black line), where each time point is modeled separately (similar to Methods 2.6b), of any defined ROI containing at least 4 active voxels from the $1^{st}$-$2^{nd}$-$6^{th}$ epoch subset GLM analysis. The standard error of the model estimate associated with each time point's LME-derived average is also included (gray shading). The blue box spanning from 0 to 20 seconds represents the 20-mmHg pressurization period. The right column depicts the corresponding ROI mask (yellow) and the activated voxels (red) contained within. The activated ROIs were as follows: **(A)** caudate putamen – 3% of mask occupied by activated voxels; **(B)** ventral dentate gyrus – 11% of mask occupied by activated voxels; **(C)** presubiculum – 29% of mask occupied by activated voxels **(D)** entorhinal cortex – 4% of mask occupied by activated voxels **(E)** septal region – 10% of mask occupied by activated voxels.

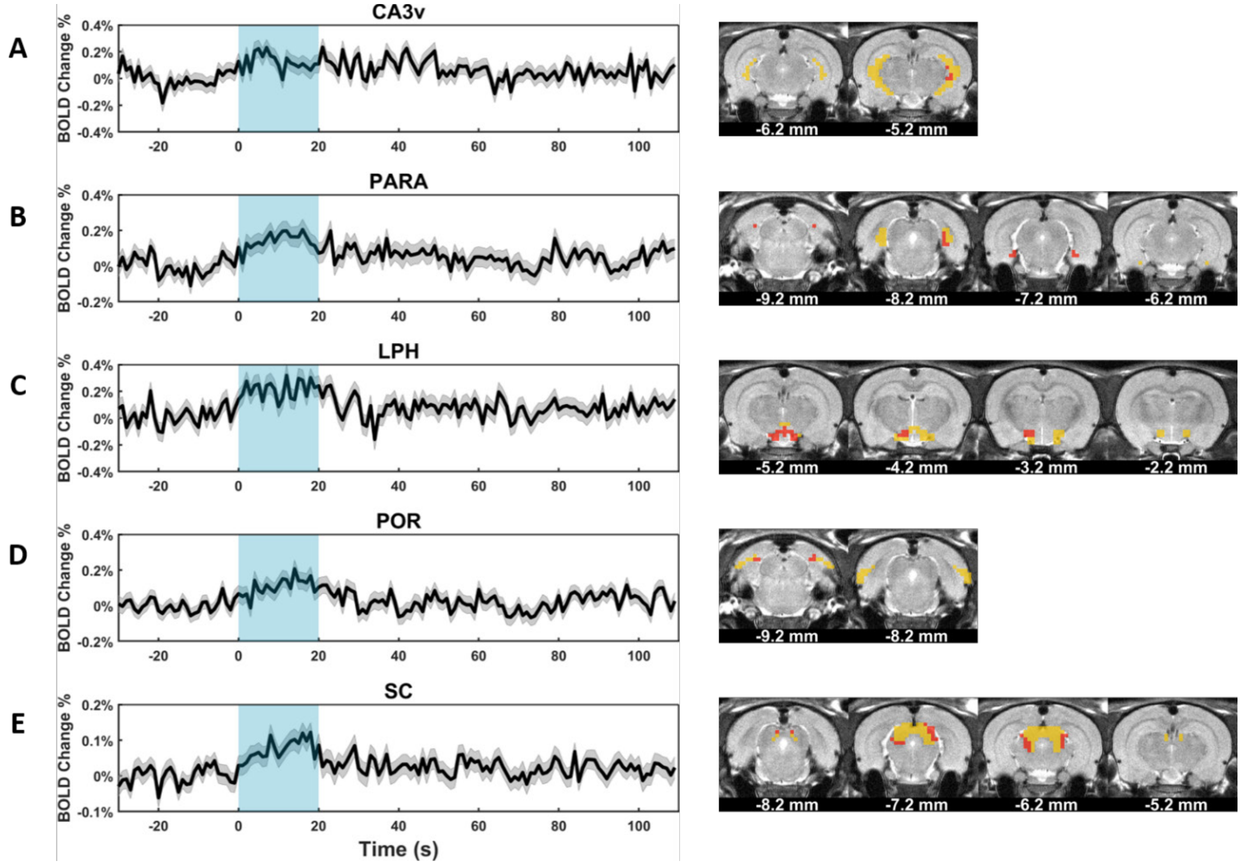


**Supplementary Figure 12: Additional GLM-derived average time courses for all active ROIs – Part II.** Panels A-E are all divided into a left and right column. The left column depicts LME-derived average time courses (black line), where each time point is modeled separately (similar to Methods 2.6b), of any defined ROI containing at least 4 active voxels from the 1$^{st}$-2$^{nd}$-6$^{th}$ epoch subset GLM analysis. The standard error of the model estimate associated with each time point's LME-derived average is also included (gray shading). The blue box spanning from 0 to 20 seconds represents the 20-mmHg pressurization period. The right column depicts the corresponding ROI mask (yellow) and the activated voxels (red) contained within. The activated ROIs were as follows: **(A)** CA3 subfield of hippocampus – 6% of mask occupied by activated voxels **(B)** parasubiculum – 39% of mask occupied by activated voxels; **(C)** lateral-posterior region of hypothalamus – 37% of mask occupied by activated voxels **(D)** postrhinal cortex – 13% of mask occupied by activated voxels **(E)** superior colliculus – 23% of mask occupied by activated voxels.

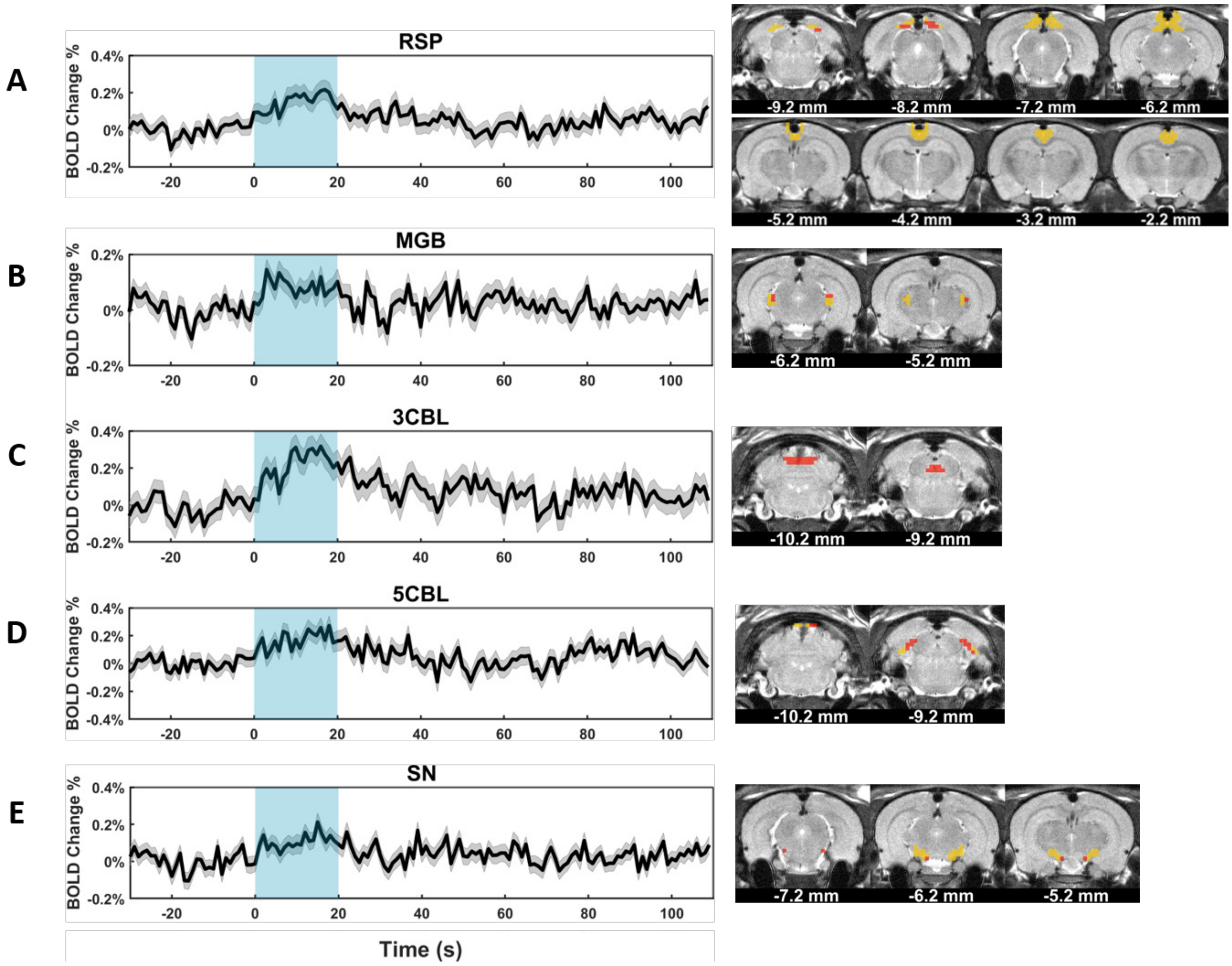


**Supplementary Figure 13: Additional GLM-derived average time courses for all active ROIs – Part III.** Panels A-E are all divided into a left and right column. The left column depicts LME-derived average time courses (black line), where each time point is modeled separately (similar to Methods 2.6b), of any defined ROI containing at least 4 active voxels from the 1$^{st}$-2$^{nd}$-6$^{th}$ epoch subset GLM analysis. The standard error of the model estimate associated with each time point's LME-derived average is also included (gray shading). The blue box spanning from 0 to 20 seconds represents the 20-mmHg pressurization period. The right column depicts the corresponding ROI mask (yellow) and the activated voxels (red) contained within. The activated ROIs were as follows: **(A)** restrosplenial area – 8% of mask occupied by activated voxels; **(B)** medial geniculate body – 25% of mask occupied by activated voxels; **(C)** 3$^{rd}$ cerebellar lobule – 100% of mask occupied by activated voxels; **(D)** 5$^{th}$ cerebellar lobule – 62% of mask occupied by activated voxels; **(E)** substantia nigra pars compacta – 14% of mask occupied by activated voxels.

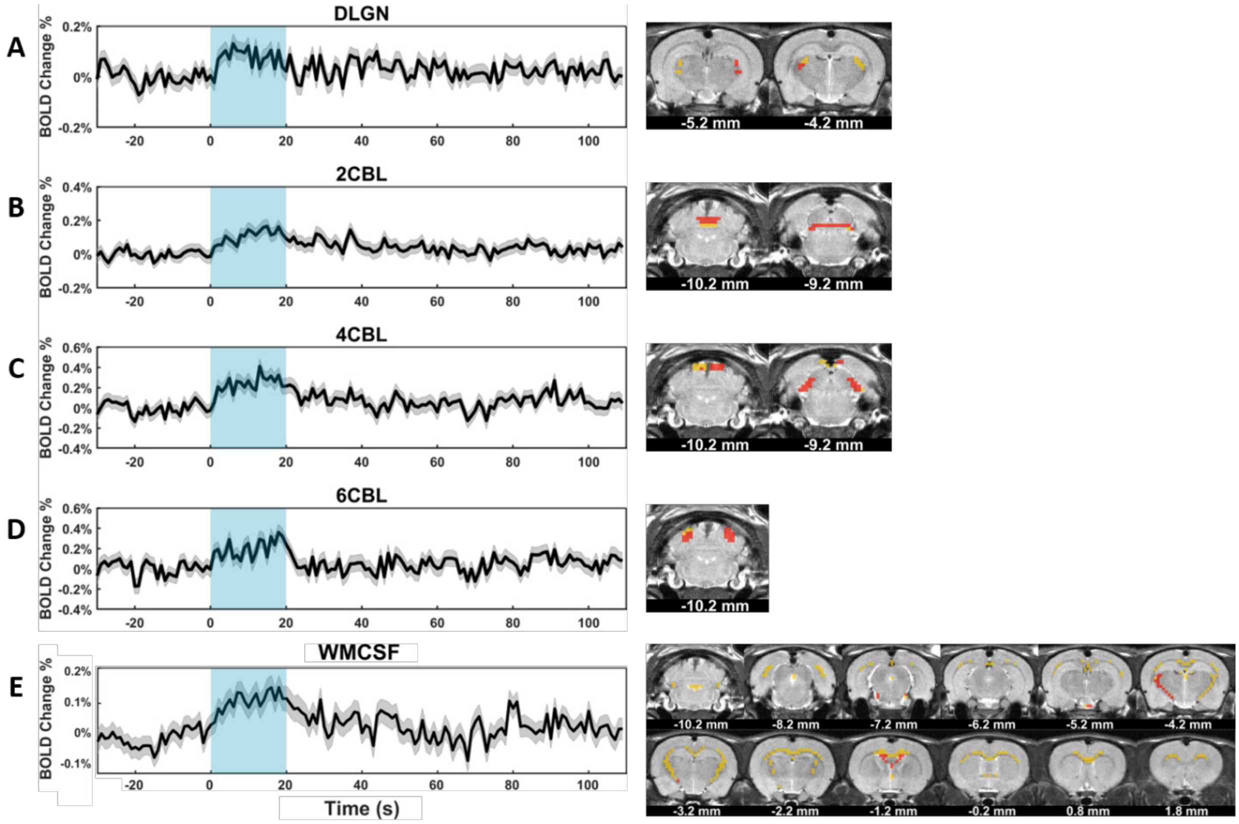


**Supplementary Figure 14: Additional GLM-derived average time courses for all active ROIs – Part IV.** Panels A-E are all divided into a left and right column. The left column depicts LME-derived average time courses (black line), where each time point is modeled separately (similar to Methods 2.6b), of any defined ROI containing at least 4 active voxels from the 1$^{st}$-2$^{nd}$-6$^{th}$ epoch subset GLM analysis. The standard error of the model estimate associated with each time point's LME-derived average is also included (gray shading). The blue box spanning from 0 to 20 seconds represents the 20-mmHg pressurization period. The right column depicts the corresponding ROI mask (yellow) and the activated voxels (red) contained within. The activated ROIs were as follows: **(A)** dorsal lateral geniculate nucleus – 35% of mask occupied by activated voxels; **(B)** 2$^{nd}$ cerebellar lobule – 82% of mask occupied by activated voxels; **(C)** 4$^{th}$ cerebellar lobule – 72% of mask occupied by activated voxels; **(D)** 6$^{th}$ cerebellar lobule – 85% of mask occupied by activated voxels; **(E)** white matter and cerebral spinal fluid – 12% of mask occupied by activated voxels

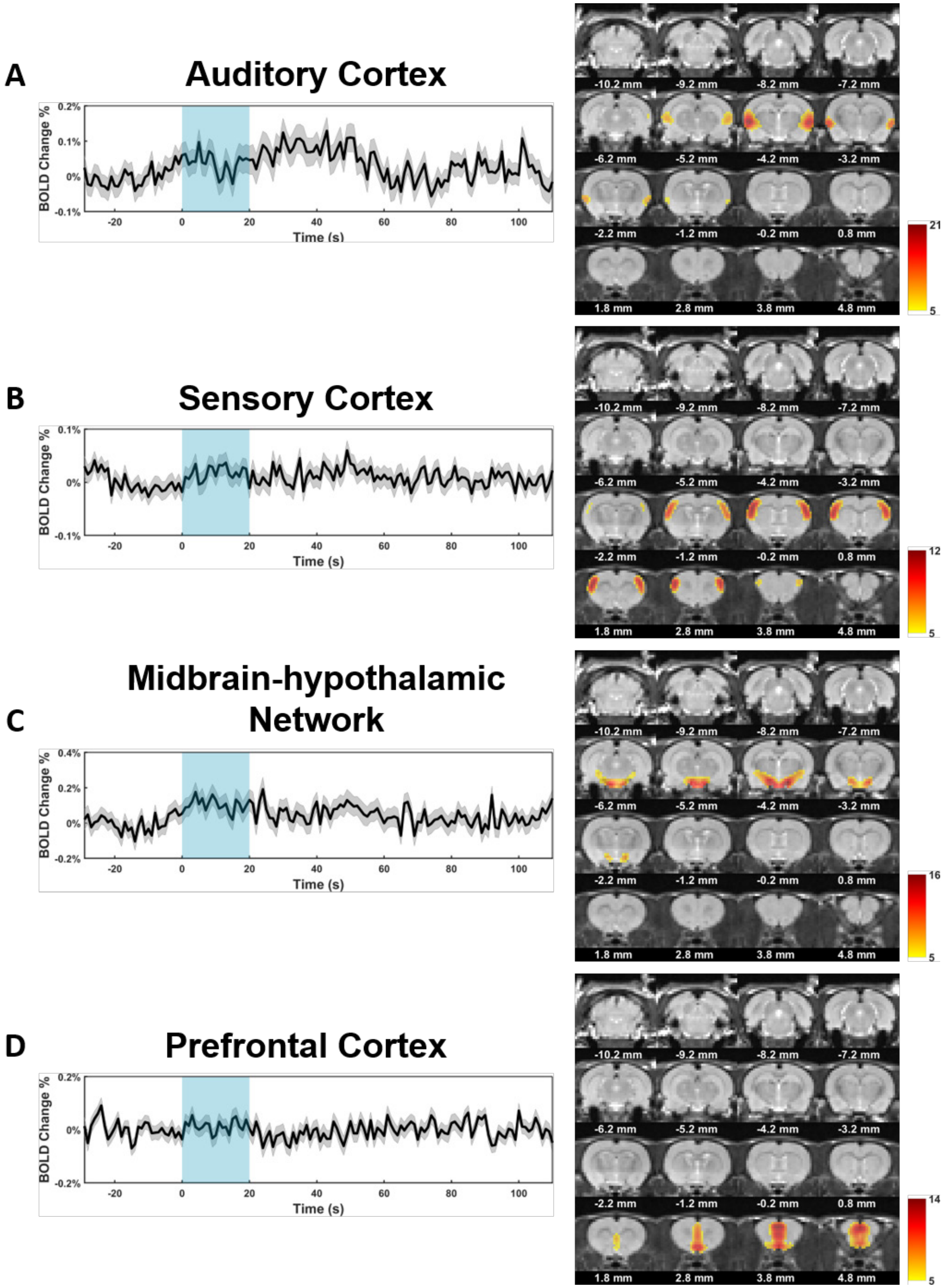


**Supplementary Figure 15: Additional sICA components derived from the 1st-2nd-6th Epoch subset – Part I.** Panels A-D are all divided into a left and right column. The left column depicts

LME-derived average time courses (black line), where each time point is modeled separately (Methods 2.6b), of spatial components calculated from the $1^{st}$-$2^{nd}$-$6^{th}$ epoch subset sICA analysis. The standard error of the model estimate associated with each time point's LME-derived average is also included (gray shading). The blue box spanning from 0 to 20 seconds represents the 20-mmHg pressurization period. The right column depicts the spatial independent component maps from which the corresponding time courses were made. In accordance with the z-value thresholding criteria, all maps contain voxels with a positive z-value ≥ 5; gray-scaled voxels of the brain have a z-value < 5. The larger the z-value of two or more voxels, the greater the degree of temporal similarity among them. Each color bar is specific to its corresponding spatial map: the lower bound (yellow) is set to z-value = 5 and the upper bound (crimson) is set to the maximum voxel-specific z-value for that particular component. The sICA component maps are best categorized as follows: **(A)** auditory cortex; **(B)** sensory cortex; **(C)** midbrain-hypothalamic network; **(D)** prefrontal cortex.

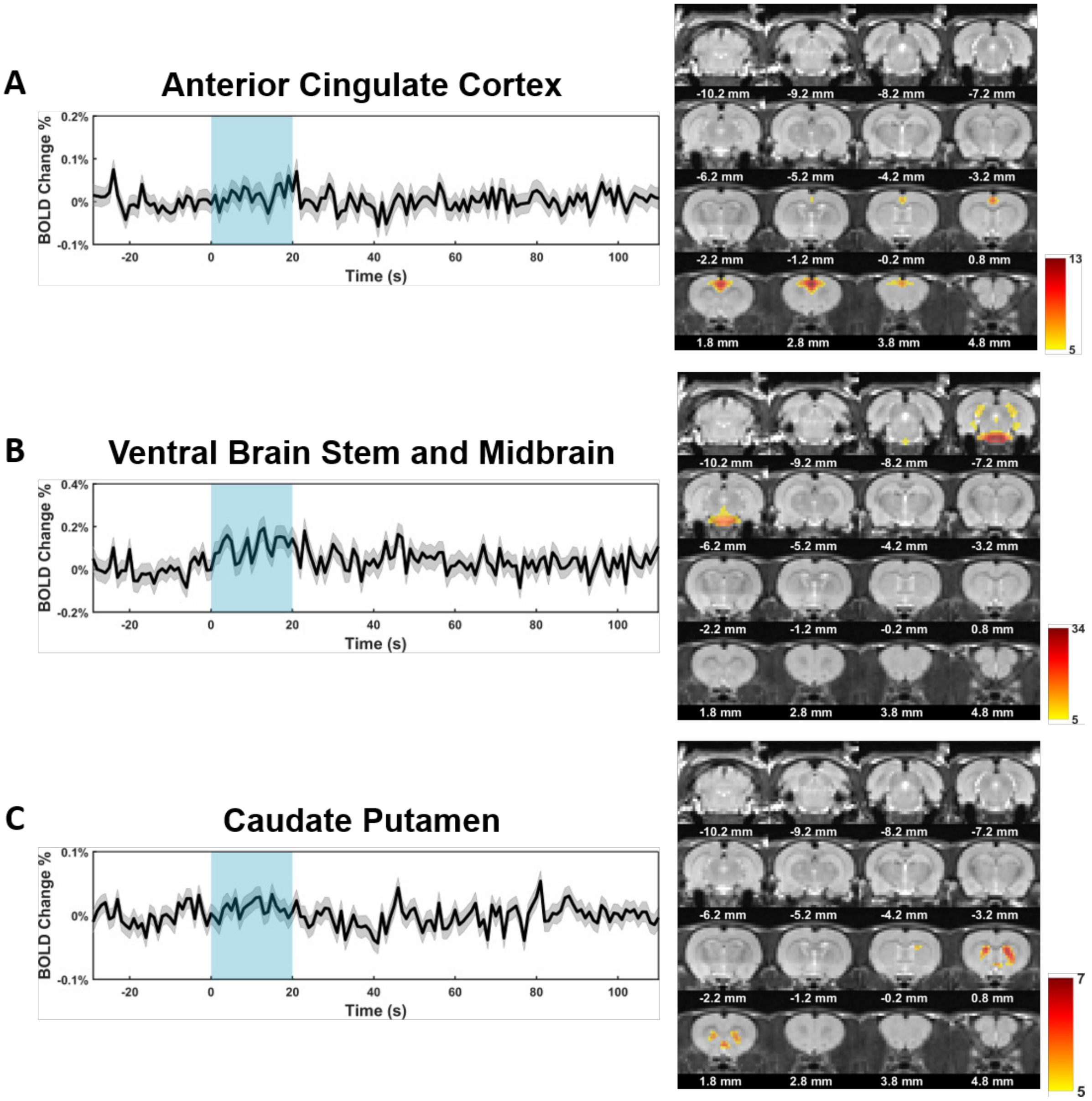


**Supplementary Figure 16: Additional sICA components derived from the $1^{st}$-$2^{nd}$-$6^{th}$ epoch subset – Part II.** Panels A-D are all divided into a left and right column. The left column depicts LME-derived average time courses (*black* line), where each time point is modeled separately (Methods 2.6b), of spatial components calculated from the $1^{st}$-$2^{nd}$-$6^{th}$ epoch subset sICA analysis. The standard error of the model estimate associated with each time point's LME-derived average is also included (*gray shading*). The blue box spanning from 0 to 20 seconds represents the 20-mmHg pressurization period. The right column depicts the spatial independent component maps from which the corresponding time courses were made. In accordance with the z-value thresholding criteria, all maps contain voxels with a positive z-value ≥ 5; gray-scaled voxels of the brain have a z-value < 5. The larger the z-value of two or more voxels, the greater the degree of temporal similarity among them. Each color bar is specific to its corresponding spatial map: the lower bound (yellow) is set to z-value = 5 and the upper bound (crimson) is set to the maximum voxel-specific z-value for that particular component. The sICA component maps are best categorized as follows: **(A)** anterior cingulate cortex; **(B)** ventral brainstem and midbrain; **(C)** caudate putamen.

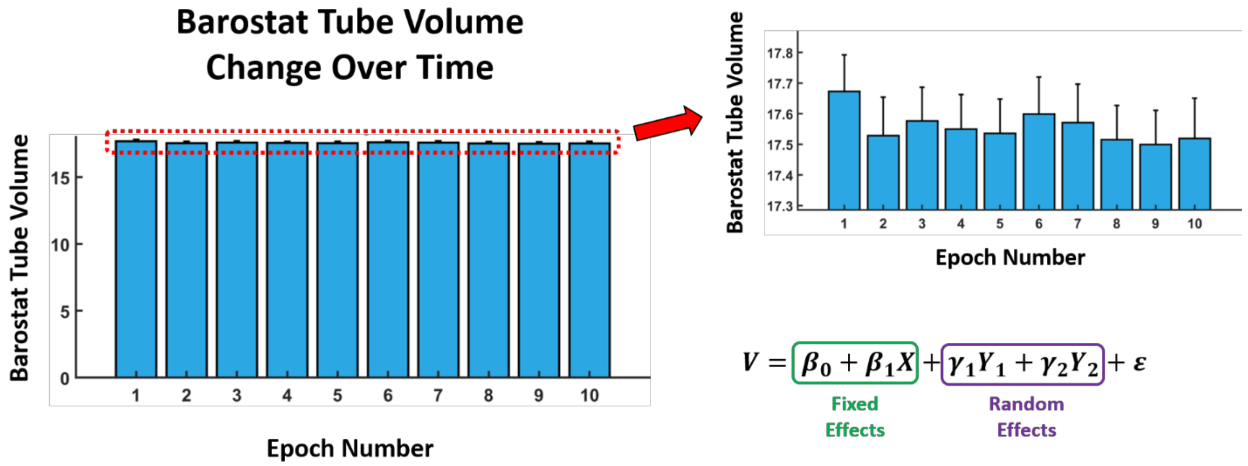


**Supplementary Figure 17: Barostat tube volume does not significantly vary as a function of epoch number.** (*Left*) A graph of barostat piston position (expressed as a tube volume) versus within-scan epoch number is shown. The bar plot height of each epoch number is determined from epoch-specific LME modeling using a similar approach as the model described in second-level GLM analysis (Methods 2.6a). The standard error of this epoch-specific model estimate is also provided. (*Top Right*) A zoomed-in version of the left graph is provided to emphasize tube volume contrast across different epoch numbers. (*Bottom Right*) The general form of the linear mixed model used to test for the presence of barostat differences across epochs is shown. Two fixed effects and two random effects were used to assess whether or not piston position was different between the 1$^{st}$-2$^{nd}$-6$^{th}$ epoch subset and the 3$^{rd}$-4$^{th}$-5$^{th}$-7$^{th}$-8$^{th}$-9$^{th}$-10$^{th}$ epoch subset (note that this is a *different* model than what was used in the bar plot representations). Fixed effects included a global average $\beta_0$ and an epoch-subset-specific coefficient $\beta_1$, where $X$ was a categorical variable taking on the value of either 0 (if the volume $V_i$ was from the first, second, or sixth epoch) or 1 (if the volume $V_i$ was from the third, fourth, fifth, seventh, eighth, ninth, or tenth epoch). The two encoded random effects were a scan-specific random intercept nested within an animal-specific random intercept. The purpose of modeling scan number as a random intercept was to account for possible differences in food content within the stomach from scan to scan (i.e. accounting for gastric emptying between scans). We failed to reject the null hypothesis of $\beta_1 = 0$ (p=0.1125). The reported $\beta_1$ estimate was 0.05518 mL.

| Summary of Subject-specific Epoch Totals | | |
|---|---|---|
| **Subject ID** | **Number of Good Epochs** | **Number of Corresponding Scans** |
| **Rat 1** | 8 | 1 |
| **Rat 2** | 17 | 2 |
| **Rat 3** | 19 | 2 |
| **Rat 4** | 19 | 2 |
| **Rat 5** | 47 | 5 |
| **Rat 6** | 17 | 2 |
| **Rat 7** | 28 | 3 |
| **Rat 8** | 77 | 8 |
| **Rat 9** | 78 | 9 |
| **Rat 10** | 86 | 9 |
| **Rat 11** | 82 | 9 |
| **Rat 12** | 74 | 8 |
| **Rat 13** | 92 | 10 |
| **Rat 14** | 55 | 6 |
| **Rat 15** | 94 | 10 |
| **Rat 16** | 97 | 10 |
| **Rat 17** | 84 | 9 |
| **Rat 18** | 97 | 10 |
| **Rat 19** | 96 | 10 |

**Supplementary Table 1: Summary of Subject-specific Epoch Totals.** This table provides a subject summary of the number of unique 'good' 140-volume epochs used throughout the analysis of this experiment. 'Good' epochs are from scans that pass the relevant motion criteria (refer to Methods 2.5). Briefly, good epochs must have FD <0.10 mm for the 28th through 53rd volume of each epoch: the 31$^{st}$ through 50$^{th}$ volume represents the distension/pressurized (20 mmHg) period of the 140-volume epoch, and the 28$^{th}$-30$^{th}$ and 51$^{st}$-53$^{rd}$ volumes represent peristimulus periods of 0 mmHg pressurization states. These peristimulus *buffers* minimize the influence of motion-induced artifacts on the gastric distension window (31$^{st}$-50$^{th}$ volumes). The total number of unique good epochs used in this paper's analysis is 1167, which belong to a total of 125 unique, 1520-volume fMRI scans.
